# Genome binning of viral entities from bulk metagenomics data

**DOI:** 10.1101/2021.07.07.451412

**Authors:** Joachim Johansen, Damian Plichta, Jakob Nybo Nissen, Marie Louise Jespersen, Shiraz A. Shah, Ling Deng, Jakob Stokholm, Hans Bisgaard, Dennis Sandris Nielsen, Søren Sørensen, Simon Rasmussen

**Author notes:** Correspondence: Simon Rasmussen.

## Abstract

Despite the accelerating number of uncultivated virus sequences discovered in metagenomics and their apparent importance for health and disease, the human gut virome and its interactions with bacteria in the gastrointestinal are not well understood. In addition, a paucity of whole-virome datasets from subjects with gastrointestinal diseases is preventing a deeper understanding of the virome’s role in disease and in gastrointestinal ecology as a whole. By combining a deep-learning based metagenomics binning algorithm with paired metagenome and metavirome datasets we developed the Phages from Metagenomics Binning (PHAMB) approach for binning thousands of viral genomes directly from bulk metagenomics data. Simultaneously our methodology enables clustering of viral genomes into accurate taxonomic viral populations. We applied this methodology on the Human Microbiome Project 2 (HMP2) cohort and recovered 6,077 HQ genomes from 1,024 viral populations and explored viral-host interactions. We show that binning can be advantageously applied to existing and future metagenomes to illuminate viral ecological dynamics with other microbiome constituents.

## Introduction

The human gut microbiota is tightly connected to human health through its massive biological ecosystem of bacteria, fungi, and viruses. This ecosystem has been profoundly investigated for discoveries that can lead to diagnostics and treatments of gastrointestinal diseases such as inflammatory bowel disease (IBD) and colon cancer as well as type 2 diabetes (T2D)^1–3^. In IBD, multiple studies have compiled a list of keystone bacterial species undergoing microbial shifts between inflamed and non-inflamed tissue sites^4,5^ and there are strong indications that the gut virome plays a role in disease etiology^6–8^. Now, the influence of bacteria-infecting viruses, known as bacteriophages, are increasingly studied and their role in controlling bacterial community dynamics in the context of gastrointestinal pathologies is slowly being unravelled^9^. Several studies have presented evidence of temperate *Caudovirales* viruses increasing in Crohn’s disease (CD) and ulcerative colitis (UC) patients^6,8,10–12^. However, it has been left unanswered if this phage expansion was due to alterations in host-bacterial abundance, thus viral-host dynamics remains another unexplored facet of the gut virome in diseases such as IBD^13^.

Today, the virome is studied through metagenomics where high-throughput sequencing is computationally processed to construct genomes of uncultivated viruses *de novo*. Viral assembly is a notoriously difficult computational task and known to produce fragmented assemblies and chimeric contigs^14^ especially for rare viruses with low and uneven sequence coverage^15,16^. For better viral assemblies, metaviromes are prepared with extra size-filtration to increase the concentration of viral particles^17,18^. However, identification of viruses without enrichment from bulk metagenomics, is increasingly utilised and overcomes the size-filtration step biases while enabling identification of primarily temperate but also lytic viruses ^19^. Currently, several approaches for identifying viral sequences in metagenomics data exist and have helped in supersizing viral databases of uncultivated viral genomes (UViGs) over the last few years ^20–22^. These tools are often based on sequence similarity^23^, sequence composition^24–27,28,29^, and identification of viral proteins or the lack of cellular ones^28,29^. A common denominator for these tools is their per-contig/sequence virus evaluation approach that is not optimal for addressing fragmented multi-contig virus assemblies.

Therefore, we developed a framework (PHAMB) based on contig binning to discover viral genome bins directly from bulk metagenomics data (MGX). For this we utilised a recently developed deep learning algorithm for metagenomic binning (VAMB)^30^ that is based on binning the entire dataset of assembled contigs. Altogether, we reconstructed 2,676 viral populations from bulk metagenomes corresponding up to 36% of the paired metavirome dataset (MVX), based on two independent datasets with paired MGX and MVX. A key development in our method is a classifier that can classify non-phage bins from any dataset with a very low error rate (3%) compared to existing virus prediction tools such as DeepVirFinder^26^ (21-65%). Our approach enables identification and reconstruction of viral genomes directly from metagenomics data at an unprecedented scale with up to 6,077 viral populations with at least one High-quality (HQ) genome by MIUViG standards^19^ in a single dataset. In addition, we show an increase of up to 210% of HQ viral genomes extracted by combining contigs into viral bins. Using this method to extract viruses from the microbial metagenomes of the HMP2 cohort we are able to delineate both viral and bacterial community structures. This allowed us to conduct an investigation of viral population dynamics in tandem with predicted microbial hosts for instance identifying 123 and 230 viral populations infecting *Faecalibacterium* and *Bacteroides* genomes, respectively.

## Results

### A framework to bin and assemble viral populations from metagenomics data

To generate the metagenomics bins we used VAMB that has the advantage of both binning microbial genomes, and grouping bins across samples into subspecies or conspecific clusters. This has proven useful for investigation of bacterial and archaeal microbiomes, but the approach has even more potential within viromics where viruses are much less conserved, more diverse, and harder to identify without universal genetic markers such as those found in bacterial organisms^31^. Clusters of conspecific viral genomes would enable straightforward identification and tracking of populations across a cohort of samples (**Figure 1a**). To develop our framework we used two Illumina shotgun sequencing based datasets with paired metagenome and metavirome available. The Copenhagen Prospective Studies on Asthma in Childhood 2010 (COPSAC) dataset consisted of 662 paired samples (^32^ and ^33^) and the Diabimmune dataset contained 112 paired samples^10^. Each of the two data sets included a list of curated viral species, 10021 and 328 respectively, that we used here as our gold standard for training and testing our tool. Compared to COPSAC, Diabimmune metaviromes had low viral enrichment (**Supplementary Figure 1**), we therefore used the average amino acid identity (AAI) model of CheckV^29^ to stratify the genomes of the metaviromes into quality tiers ranging from Complete, High Quality (HQ), Medium Quality (MQ), Low Quality (LQ), and Non Determined (ND) to establish a comparable viral truth.

**Figure 1.**
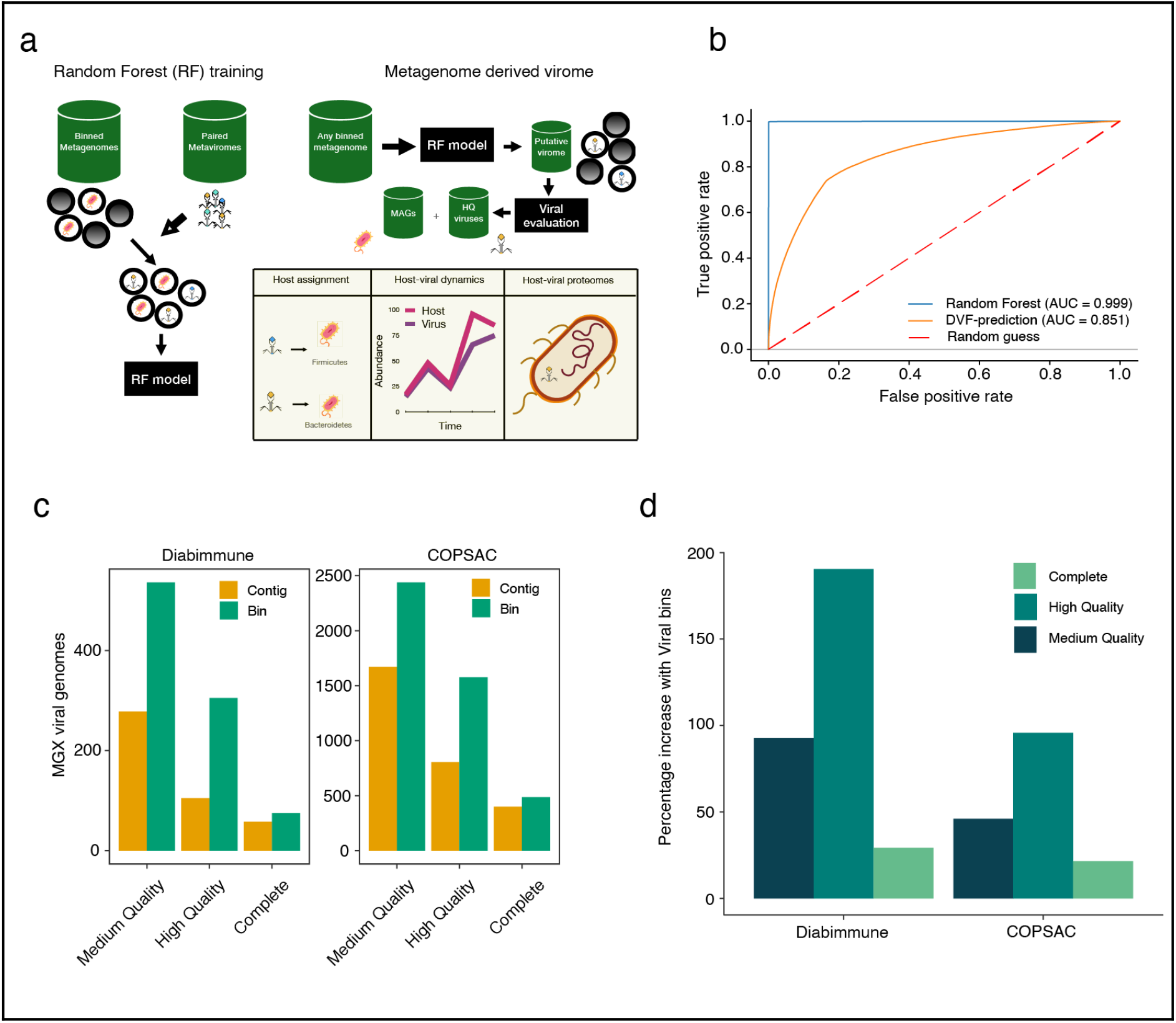
A framework to bin and assemble viral populations from metagenomics data. **a)** Illustration of workflow to explore viruses from binned metagenomes. First the RF model was trained on binned metagenomes; bacterial bins were identified using reference database tools and viruses were identified using assembled viruses from paired metaviromes. Viral and bacterial labelled bins were used as input for training and evaluating the RF model. Bins from any metagenome can be parsed through the RF model to extract a space of putative viral bins that are further validated for HQ viruses using dedicated tools like CheckV. Binned MAGs and viruses can then be associated in a host assignment step. Host-viral dynamics can be explored in longitudinal datasets to establish temperate phages and the contribution of viruses to Host pangenomes **b)** AUC performance curves for the Random Forest model and using *de novo* viral prediction method for annotating viral bins. Predictions were produced based on bins annotated as either Viral or Bacterial. **c)** The number of viral genomes recovered from bulk metagenomes, counted at three different levels of completeness in Diabimmune or COPSAC cohorts, evaluated as either single-contigs or viral bins from bulk metagenomes. Evaluation of genome completeness was determined using CheckV here shown for MQ >=50%, HQ >=90%, Complete =100%). Closed genomes are annotated as “Complete” based on direct terminal repeats or inverted terminal repeats. **d)** The percentage-increase of viral genomes found in Diabimmune or COPSAC cohorts using our approach relative to single-contig evaluation. The increase is coloured at three different levels of completeness determined using CheckV, corresponding to the ones used in (c). Abbreviations: MAGs = Metagenome-assembled genomes, HQ: High-quality, MQ: Medium-quality. AUC: Area under curve.

### Viral binning is more powerful compared to single-contig approaches

The output of binning metagenomics samples can be hundreds of thousands of bins and we therefore first developed a Random Forest (RF) model to distinguish viral-like from bacterial-like genome bins. The RF model takes advantage of the cluster information from binning and aggregates information across sample-specific bins to form subspecies clusters. Here, we found that the RF model was able to separate bacterial and viral clusters very effectively with an Area Under the Curve (AUC) of 0.99 and a Matthews Correlation Coefficient (MCC) of 0.98 on the validation set (**Figure 1b** and **Supplementary Table 1**). Compared to single-contig-evaluation methods such as DeepVirFinder, our approach was far superior as DeepVirFinder only achieved an AUC of 0.85 and MCC of 0.06. This difference in performance is likely explained by the RF model evaluating on both cluster and bin-level where one sequence with a low viral score does not lead to a misprediction of the whole bin. For instance, we achieved an increase of 200 (190%) and 771 (95%) HQ bins recovered for the Diabimmune and COPSAC datasets compared to using single-contig-evaluation according to CheckV **(Figure 1c-d)**. In addition, we observed a significantly greater number of viral hallmark genes per virus when using viral bins in both datasets (Two-sided T-test, P<0.0005), while the length and viral fraction were largely comparable (**Supplementary Figure 2)**.

### Binning the metagenome identifies viral genomes not identified from the metavirome

When applying our method of binning with VAMB and the RF model we obtained 4,480 and 916 viral bins with a MQ or HQ representative bin across the COPSAC and Diabimmune datasets, respectively. We then considered all VAMB clusters as ‘viral populations’ and thus obtained 2,428 and 534 viral populations with at least 1 MQ or better viral bin. After comparing the viral populations obtained from the metagenomics datasets to the respective metaviromes we recovered 17-36% of HQ viruses (corresponding to 527 and 2,676 metaviromic viral populations) established in the metaviromes on species (ANI>95) level and 9-28% on strain (ANI>97) level (**Figure 2a**). The fraction of viruses in the metavirome recovered in the metagenome was considerably higher than more recent estimates^34^, which estimated 8.5-10%. This was interesting since the deeply sequenced metavirome may capture multiple low abundant viruses typically not found in metagenomes. Additionally, we found that 46-69% of the HQ metagenome viral populations, corresponding to 124 in Diabimmune and 839 viral populations in COPSAC, were not found in the metavirome, suggesting that a significant part of the virome may be lost during viral enrichment or not represented in induced forms as they are integrated prophages (**Figure 2b**). However, we also found that 65-83% of the HQ viral populations in the metavirome were not found in the metagenome data (total 197 in Diabimmune and 2,589 in COPSAC) suggesting the reverse to be true as well. For a subset of the viruses found in the COPSAC bulk and metavirome, we estimated a higher mean completeness with viral bins (Two-sided T-test, P=2.2e-16) (**Figure 2c**).

**Figure 2.**
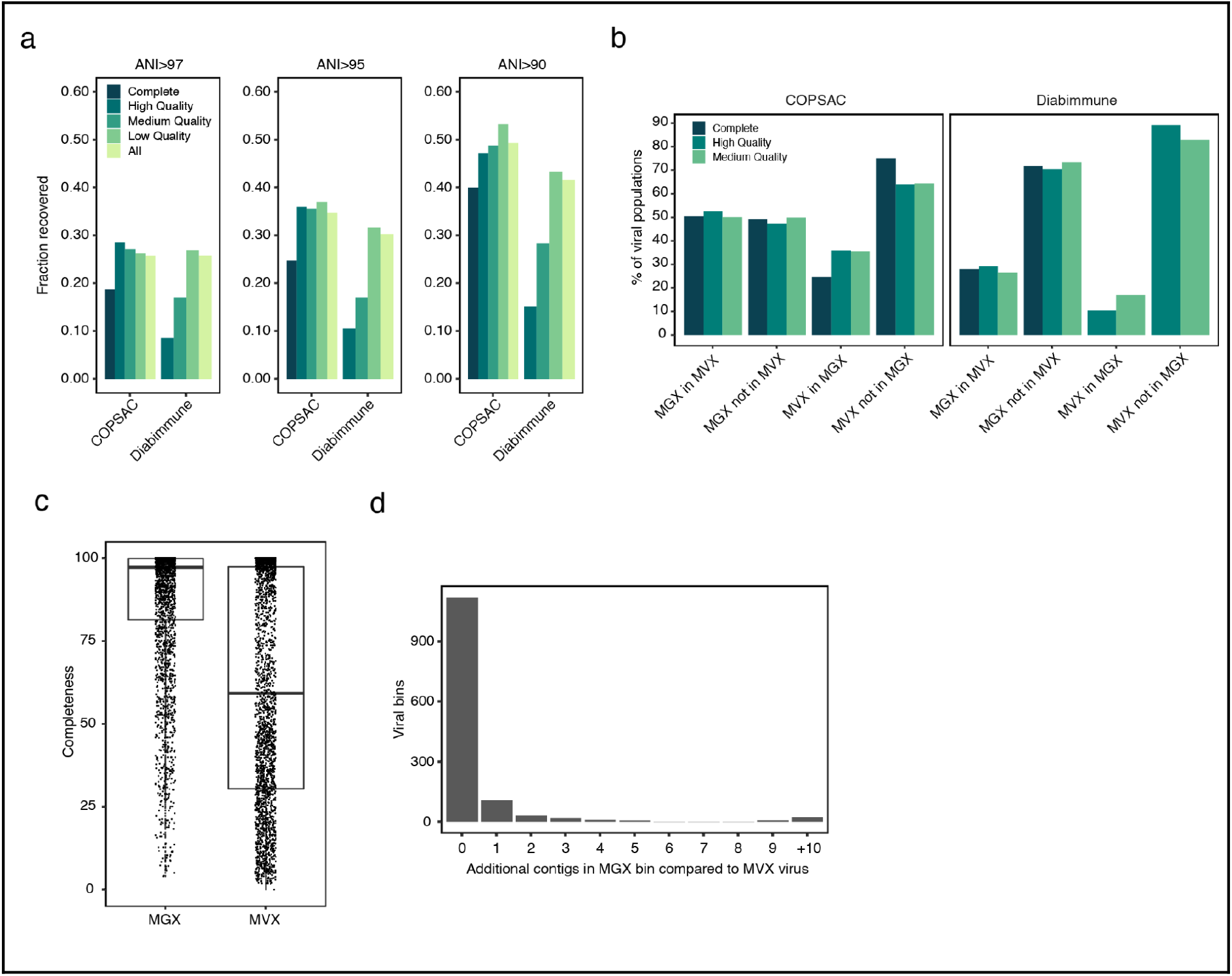
Binning the metagenome identifies viral genomes not identified from the metavirome. **a)** The fraction of metavirome viruses in COPSAC and Diabimmune coloured at different levels of completeness or all together determined with CheckV, identified in VAMB bins from bulk metagenomics of the same cohorts. We defined a metavirome virus to be recovered if the aligned fraction was at least 75% and ANI was either >90,>95,>97.5 to a VAMB bin based on FastANI. **b)** The percentage of viral populations, at different levels of completeness determined with CheckV, identified in both metaviromes (MVX) and bulk metagenomics (MGX) or unique to either dataset. Shared populations are identified with a minimum sequence coverage of 75% and ANI above 95%. (1) MGX in MVX: % of Viral populations found in MGX also found in MVX. (2) MGX not in MVX: % of Viral populations unique to MGX i.e. not found in MVX. (3) MVX in MGX: % of Viral populations found in MVX also found in MGX. (4) MVX not in MGX: % of Viral populations unique to MVX i.e. not found in MGX. **c)** Viral genome completeness estimated for 2,646 viruses found both in metaviromes and bulk metagenomics sharing the same nearest reference in the CheckV database. **d)** The number of contigs in Viral bins from bulk metagenomics that do not align to the closest viral reference in the metavirome. In the majority of viral bins all contigs align to the nearest reference. ANI: Average nucleotide identity %.

Altogether we found that a great proportion of the gut viral populations can be reconstructed from the metagenomics data and retrieved with even higher completeness compared to the metavirome counterparts.

### Viral bins have low contamination

Lastly, we wanted to investigate the occurrence of technically “misbinned” and contaminating contigs that could inflate viral genome size and influence evaluation and other downstream analyses. Based on 1,340 viral bins highly similar to metavirome viruses in COPSAC (see methods), we found in 83.7% of all cases, every contig in the bin mapped to the virus (**Figure 2d)**. In total 8.1% of the viral bins contained more than one contig not mapping to the corresponding metavirome-virus indicating contamination, variation, or incompleteness. In summary, our combined binning and machine learning approach improves identification and recovery of viral genomes from metagenomics data and outlines the possibility of binning fragmented viruses directly from human gut microbiome samples.

### Reconstructing the virome of a the HMP2 IBD gut metagenomics cohort

We then applied our method to the HMP2 IBD cohort consisting of 27 healthy controls, 65 CD, and 38 UC patients^35^. These samples were gathered in a longitudinal approach and consisted of between 1-26 samples per patient. Importantly, no characterised metaviromics data is available from this cohort and using our approach we were able to identify bacterial and viral populations in the cohort and explore their dynamics in IBD using only metagenomics data. From the cohort we recovered 577 Complete, 6,077 HQ, 9,704 MQ (**Figure 3a)**, and 122,107 LQ viral bins corresponding to 263 Complete, 1,024 HQ, 2,238 MQ, and 44,017 LQ viral populations. We also observed an increase in genome completeness for larger viruses/jumbo viruses with a genome size > 200 kbp^36^ compared to single contig evaluation (**Supplementary Figure 3)**. In addition, we observed that similar viral length distributions for viruses recovered as a single contig and as viral bins, both correlated with CheckV quality tiers **(Figure 3b)**.

**Figure 3.**
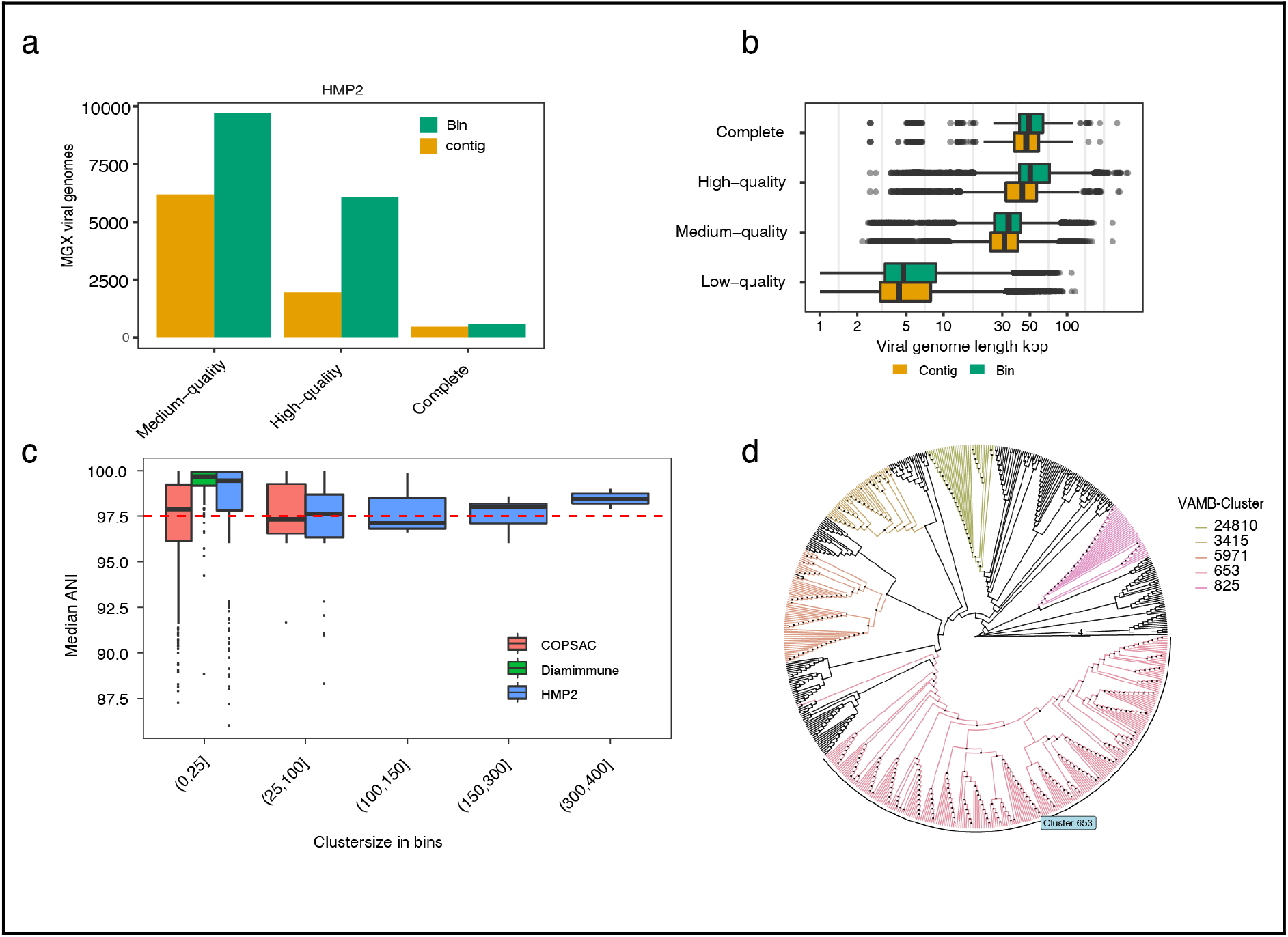
Reconstructing the virome of a human gut metagenomics cohort. **a)** The number of viral genomes with 3 different levels of completeness in HMP2, evaluated as either single-contigs or viral-bins from bulk metagenomes. Evaluation of genome completeness was determined using CheckV here shown for Medium-quality >=50% (MQ), High-quality >=90% (HQ), Complete =100%. Closed genomes are annotated as “Complete” based on the presence of either DTR or ITR. **b)** The sequence length distribution in kbp of viral genomes at 4 different levels of completeness in HMP2, evaluated as either single-contigs or viral-bins from bulk metagenomes. Shown for Low-quality <50%, MQ >=50%, HQ >=90% or Complete =100%. Closed genomes are annotated as “Complete” based on the presence of either DTR or ITR. **c)** Median ANI based on pairwise ANI genome measurements between bins within the same VAMB cluster. Median ANI is consistently above 97.5 in small VAMB clusters with 0-25 bins and in larger VAMB clusters with 300-400 bins. **d)** Cladogram of an unrooted phylogenetic tree with crAss-like bins based on the large terminase subunit protein (TerL). Five different VAMB clusters have been coloured and illustrate high monophyletic relationships. The phylogenetic tree was constructed using IQtree using the substitution model VT+F+G4. ANI: Average nucleotide identity %. DTR: Direct terminal repeats. ITR: Inverted terminal repeats. Kbp: Kilo base pairs.

### Viral population taxonomy is highly consistent

We then investigated the taxonomic consistency of our viral populations and found this to be very high as the median intra-cluster Average Nucleotide Identity (ANI) for MQ to Complete viral clusters was 97.3-99.3% **(Supplementary Figure 5)**. Even in clusters with over 100 sample-specific viral bins the intra-cluster median ANI was consistently high (median=97.1-98.5%) **(Figure 3c).** Inter-cluster ANI was much lower in the 91.7-92.8% range closer to genus level. Therefore, our approach was able to identify and cluster near strain-level viral genomes across samples. For example, in the HMP2 dataset, we identified 50 different viral populations for a total of 916 MQ or better crAss-like viral bins. Here, viral population 653 corresponded to the prototypic crassphage^37^ and accounted for 253 of the 916 crAss-like genomes discovered in the HMP2 dataset. We then used all of these 916 bins to generate a phylogenetic tree based on the large terminase subunit (TerL) and found highly consistent placement of the viral genomes according to their binned viral population **(Figure 3d, Supplementary Figure 6)**. Viral population 653 formed one monophyletic clade except for one bin while all the other crAss-like clusters were monophyletic. The division of the crAss-like genomes into the binned clusters therefore likely represents actual viral diversity. Taken together, this shows that our reference-free binning produces taxonomically accurate viral clusters, thus aggregating highly similar viral genomes across samples.

### The metagenomic virome is personal and highly stable in healthy subjects

Several metavirome studies have reported the presence of stable, prevalent, and abundant viruses in the human gut^7,38^. We found that the gut virome in the HMP2 cohort^35^ was highly personal and stable over time in nonIBD subjects, which was reflected by the lower Bray-Curtis dissimilarity between samples from nonIBD subjects compared to UC (T-test, P=0.015, CI= −0.01;−0.13) and CD subjects (T-test, P=0.017, CI = −0.12;−0.01) (**Figure 4a-b).** In addition, the dysbiotic samples, as defined by Price *et al.* (2019)^35^, could be clearly separated with a principal component analysis (PCoA), where the virome explained 4.2% and 3.4% of variation (**Figure 4c).** This was confirmed with a permanova-test on viral (P<10−3, R^2^=1.6%) and bacterial abundance profiles (P<10−3, R^2^=3.0%) and shows dysbiosis affecting both the virome and bacteriome. Alpha-diversity metrics supported this as Shannon Diversity (SD) was higher in nonIBD subjects compared to both UC and CD (T-test, P=4.80e-06 and P=2.74e-09) while dysbiosis affected every patient group resulting in a significantly reduced SD. In accordance, viral richness was lower in UC (T-test, P=<2e-16, CI=−12.40;−19.80) and CD (T-test, P=<2e-16, CI=−12.91;−19.50) patients and further exaggerated in dysbiotic samples **(Figure 4d+e)**. These viral alpha-diversity trends were also observed in the bacteriome,suggesting that the viruses follow the expansion or depletion of their bacterial host during dysbiosis (**Supplementary Figure 8)**. Indeed, we identified 250 likely temperate viruses out of 348 differentially abundant viruses that expanded with increasing dysbiosis (linear-mixed effect model, adj. P < 0.005, FDR-corrected). This observation acknowledges earlier results showing an increase in temperate viruses in UC and CD^6,11^. Further analysis on the longitudinal abundance profiles of virus and predicted bacterial host reaffirmed the synchronised expansion theory (**Supplementary Figure 9)**.

**Figure 4.**
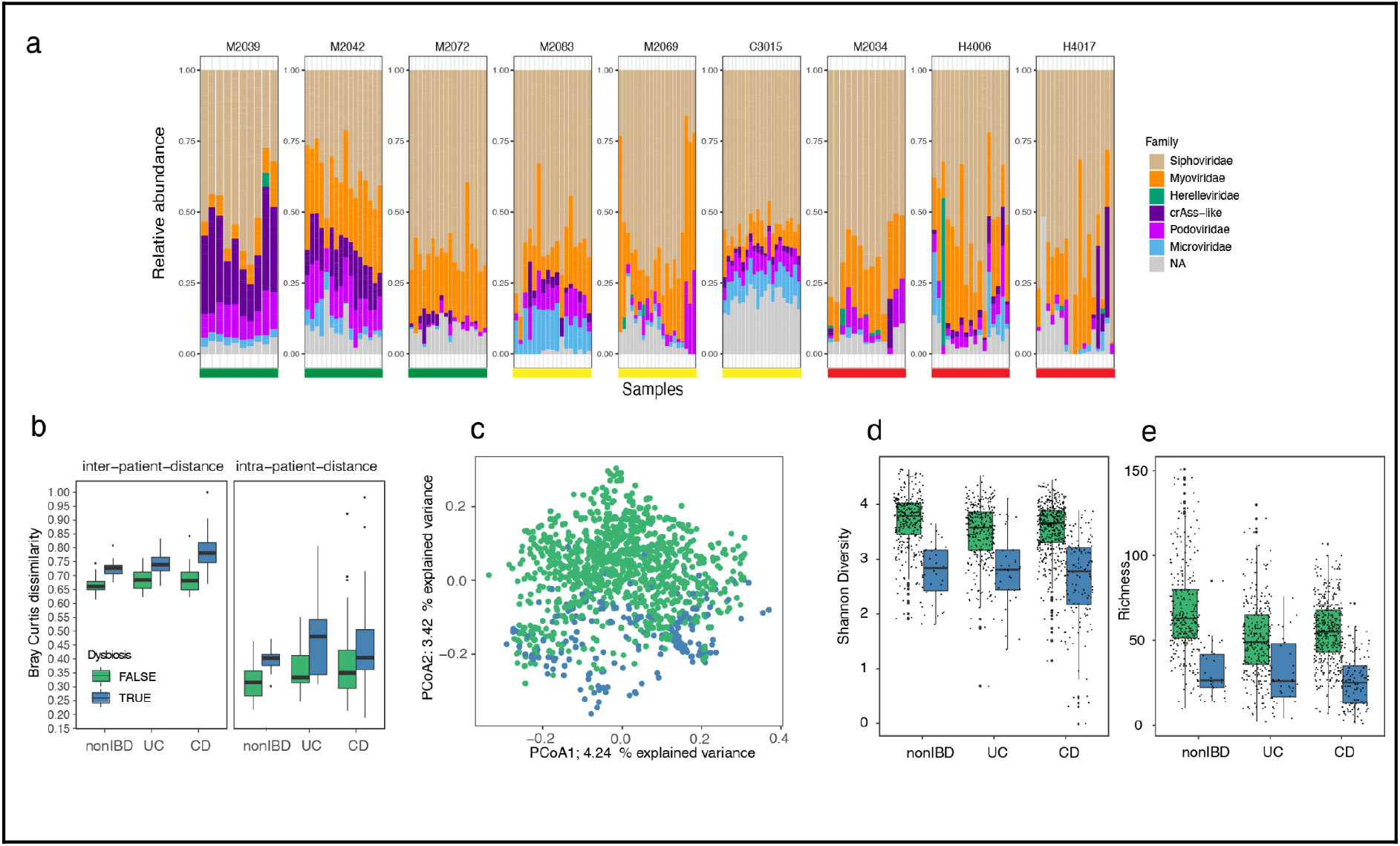
The metagenomics estimated virome is personal and highly stable in healthy controls. **a)** Longitudinal virome compositions for 3 non-IBD (green bar), 3 UC (yellow bar) and 3 CD (red bar) diagnosed subjects. Each panel represents a subject where the virome composition is organised according to the total relative abundance according to the taxonomic viral family, where “NA” populations are coloured grey. **b)** Dissimilarity boxplots based on Bray Curtis distance (BC) function between samples from different subjects (first panel inter-patient-distance) and between samples from the same subject (second panel intra-patient-distance). The BC distances are shown for samples from non-IBD, UC and CD diagnosed subjects. Furthermore, BC distances are coloured according to dysbiosis (blue) or not (green). **c)** Principal component analysis (PCoA) of Bray-curtis distance matrix calculated from the viral abundance matrix in HMP2. Each point is coloured according to diagnosed dysbiosis as in b. **d)** Shannon-diversity estimates of metagenomics derived viral populations and coloured according to dysbiosis as in b. **e)** Per sample viral population richness based on the number of viral populations detected (abundance > 0) in the samples. Coloured according to dysbiosis as in (b). non-IBD: healthy control, UC: Ulcerative colitis, CD: Crohn’s disease.

### Viral-host interactions can be explored from viral populations and MAGs

A unique feature of performing the analysis on metagenomics data is that both the bacterial and viral populations are binned simultaneously. Therefore, we are able to estimate abundance of both the viral and bacterial compartment of the microbiome and explore the viral host range *in silico* using the MAGs. In total from the HMP2 dataset, we obtained 3,130 and 3,819 Near Complete (NC) and Medium Quality (MQ) MAGs^39^. Based on MAG-derived CRISPR spacers we found spacer-hits to 464 (45.3%) to viral populations with at least one HQ representative. To further expand our viral-host prediction we conducted an all-vs-all alignment search between the MAGs and viral populations for prophage signatures. Then by combining the CRISPR spacer and prophage search we connected 93.6%, 74.4%, 82.5%, and 65.0% of MAGs from *Bacteroidetes*, *Firmicutes, Actinobacteria*, and *Proteobacteria* phylum, respectively, with at least 1 virus **(Supplementary Figure 10)**. We estimated host-prediction purities to be 94.5% and 75.6% on species rank for the CRISPR-spacer and prophage signature (**Supplementary Figure 11B**). Therefore, we confirmed that most gut phages have a primarily narrow host-range^40^. MAGs belonging to the genera *Faecalibacterium* and *Bacteroides* seemed to be viral hotspots since 99.7% to 98.7% could be associated with a HQ viral bin, corresponding to 123 and 230 distinct viral populations, respectively (**Figure 5a+b)**. For instance, in abundant commensals like *Bacteroides vulgatus* (cluster 216) we observed consistent prophage signals over time for multiple viruses across several samples (**Figure 5c)**. Interestingly, because the host range of crAss phages are not well understood we investigated CRISPR spacer hits to the MAGs in our databases. Even though we could host-annotate an overall of 45.3% of all HQ viral populations to a MAG, none of the 916 crAss-like bins could be associated with any of the 3,306 Bacteroidetes bins in our dataset, which suggests that crAss-like phages are not frequently targeted by CRISPR spacers extracted from Bacteroidetes CRISPR-Cas systems.

**Figure 5.**
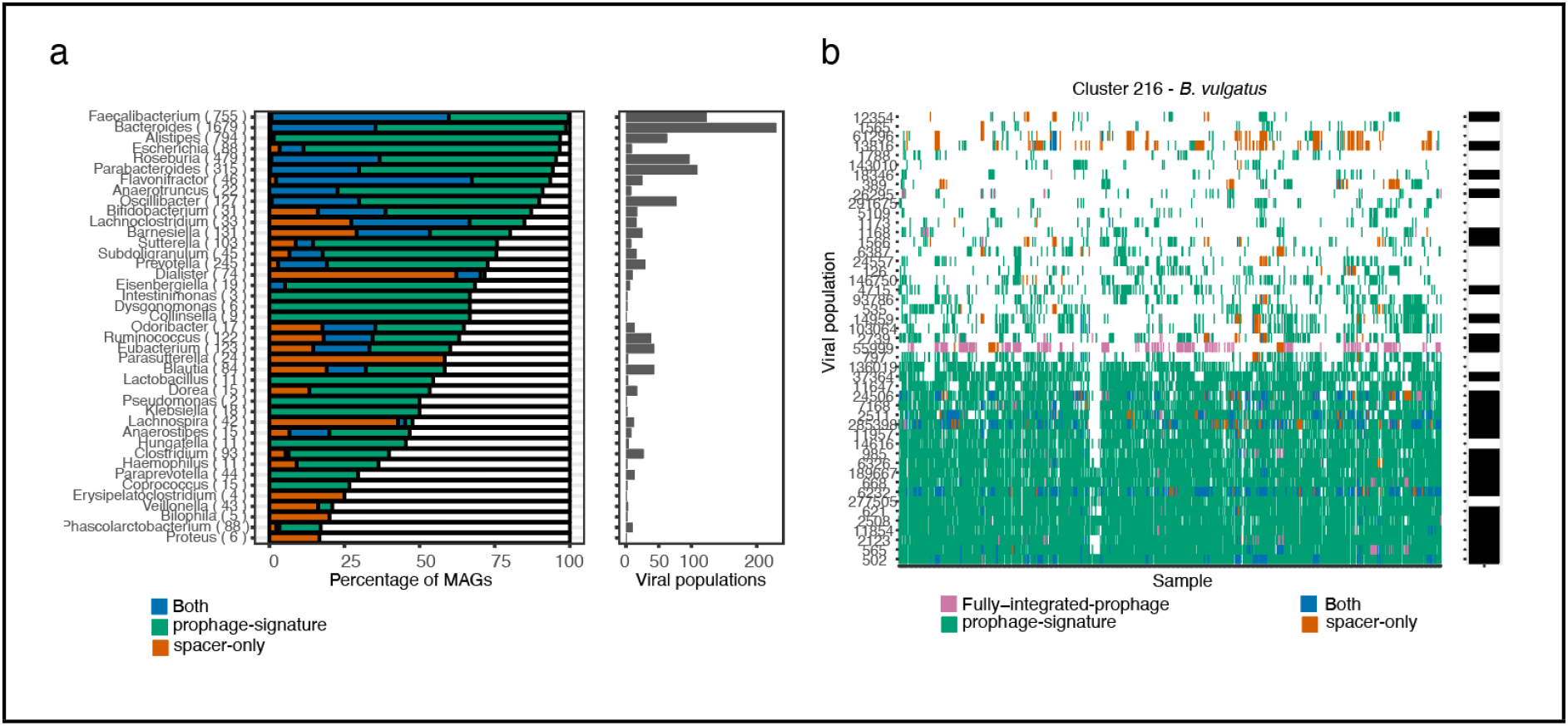
Viral-host interactions can be explored from viral populations and MAGs. **a)** Bacterial MAGs and viral relations. Each MAG was connected to the viral bins using either sequence alignment of virus to MAG (green), CRISPR spacer alignment (orange) or both (blue). The right panel shows the percentage of MAGs, grouped by genera, that was annotated with the virus via alignment og CRISPR spacer. **b)** The number of distinct viral populations associated with a MAG genera based on either of the following: sequence alignment of virus to a MAG within the given genera, CRISPR spacer alignment or both. **c)** Viral association to all MAGs of VAMB cluster 216 (*Bacteroides vulgatus*) in the HMP2 dataset. For instance, viral population 502 was associated with the *B. vulgatus* across the vast majority of samples where *B. vulgatus* was present.

### The binned viral populations are enriched in proteins found in temperate phages

Another topic of interest was viral-host complementarity, in particular what functions bacteriophages could provide to the host and how the viral proteome differs with respect to host taxonomy. Using our map of viral-host connections and through characterization of viral protein sequences we ranked protein annotations stratified by their predicted host genera. Overall, the proteins were highly enriched for annotations related to viral structural proteins such as baseplate, portal, capsid, head, tail/tail-fibre, and tail tape measure but also viral integrase enzymes and Lambda-repressor proteins (**Supplementary Table 4**). For instance, Lambda-repressor proteins were found in up to ~60% of all viruses infecting *Faecalibacterium* suggesting that our dataset is enriched with temperate phages (**Figure 6a)**. Interestingly, we also identified virally encoded proteins domains, which are known to function as viral entry receptors^41^, to be enriched within a group of viral populations infecting *Bacteroides* and *Alistipes* such as the TonB plug and TonB dependent receptor domains (PF07715 and PF00593, Fisher’s exact test, adj. P <0.005, FDR-corrected) and an outer membrane efflux protein also known as TolC (PF02321, Fisher’s.exact, adj. P <0.005, FDR-corrected) (**Supplementary Table 5**). Furthermore, the TonB domains also encode an established immunodominant epitope^42^ suggesting that viral populations carry immunogenic entry receptors when expressed by their host. Finally, Reverse Transcriptase (RT, PF00078) proteins were also highly detected, in agreement with recent results^21^ and shared by all viral populations irrespective of the predicted host (**Supplementary Figure 12A)**.

**Figure 6.**
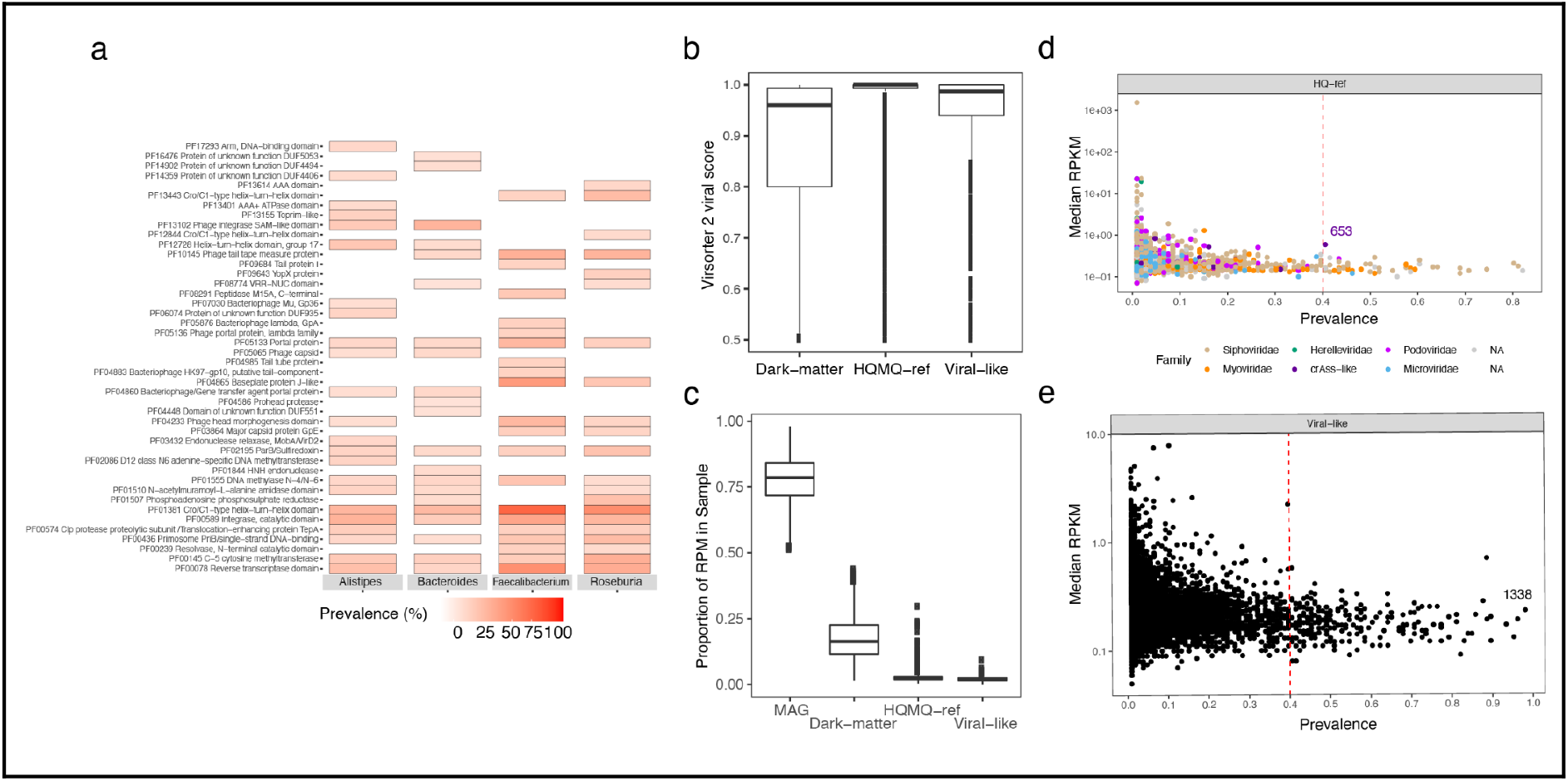
Viral proteins and the dark matter meta-virome. **a)** The percentage of HQ viruses, associated with four bacterial host genera; Alistipes, Bacteroides, Faecalibacterium and Roseburia, which encode top-20 prevalent PFAM domains. **b)** Virsorter2 viral prediction scores for all viral bins with at least one viral hallmark gene. Completeness was estimated using CheckV and the bins were grouped as (1) HQMQ-ref when completeness >=50% or High-quality >=90%, (2) bins with less than 50% completeness were annotated as Dark-matter, and (3) dark-matter bins with confident CRISPR-spacers against a bacterial host were annotated as Viral-like. **c)** The distribution of sample RPM of bacterial MAGs, HQMQ-ref viral populations, Dark-matter and Viral-like populations as defined in (b). The majority of sample reads were mapped to MAGs but on average 17.7% of all reads mapped to Dark-matter bins. **d)** The abundance in RPKM of rare and highly prevalent viruses with a HQ genome in HMP2. Each point represents a viral population coloured according to viral taxonomic family. The progenitor crassphage is indicated as cluster 653. **(e)** As in (d) but with Viral-like populations like cluster 1338 showing that many are low abundant, but highly prevalent. Abbreviations: RPM: Read per million. RPKM: Read per kilobase million.

### Exploring the dark matter metavirome

Finally, we investigated the part of the RF predicted bins that did not resemble any of the known genomes, i.e. metagenomics “dark-matter”. These were defined as populations without at least one HQ or MQ viral bin. Such populations therefore represent a part of the microbiome that are not classified as bacterial, archaeal and not alike known viral genomes. Since dark-matter populations were numerous (97.6% of all RF predicted VAMB clusters) we suspected many of these to be fragmented viruses or unknown viruses. Dark-matter populations larger than 10 kbp with at least 1 viral hallmark gene displayed lower viral prediction scores compared to HQ-MQ viral bins, while bins targeted by CRISPR-spacers displayed a significantly higher prediction score (T-test, CI=0.05:0.067, P = 2.2e-16), thus we annotated these as “viral-like” (**Figure 6b, Supplementary Figure 13**). When stratifying read abundance on these groups (HQ-MQ, viral-like, dark-matter) we found them to explain on average 2.77%, 2.04%, and 17.7% of total read abundance across samples, respectively (**Figure 6c**). Furthermore, we found that 5% HQ and 3.7% viral-like populations were detected in at least 40% of the patients across disease states. For instance, HQ viral populations cluster 653 were observed in 41% of the cohort, respectively (**Figure 6d)**. Simultaneously, viral-like population 1338 was observed in 98% of individuals but displayed a low similarity to any reference genome (**Figure 6e)**. Evidently, a significant portion of the sequenced microbiome remains dark-matter while HQ viruses identified in this study only accounts for a small fraction of the sequenced space.

## Discussion

Because of the current challenges facing the viral assembly process, which results in partial and fragmented viral genome recovery^14,16^, the viral domain of life has traditionally been notoriously difficult to study. Metavirome datasets have been crucial for identifying a broad scope of viruses, in particular virulent ones. However, the paucity and difficulties in creating metavirome datasets combined with the fact that bulk metagenomes are produced in abundance, calls for more methods to efficiently extract the viromes found therein. Here we present an improved framework for exploring metavirome directly from bulk metagenomics datasets.

Using our map of viral and bacterial connections we wanted to associate and study the human gut virome in sync with highly abundant gut bacteria such as *Bacteroides* and *Faecalibacterium*. Several of these genera represent not only highly abundant gut commensals but also hotspots for viruses as we have shown by connecting 230 and 123 viral populations to *Bacteroides* and *Faecalibacterium*, respectively. In agreement with other results^12^, we found that *F. prausnitzii* genomes are rich in prophages and were able to annotate one for 99.7% of the bacterial bins in HMP2. In the HMP2 cohort we identified 250 likely temperate *Caudovirales* viruses expanding in a synchronised manner with bacterial hosts following increasing gut dysbiosis^6,11^. However, more work is needed to outline the intricate virus-host dynamics that can explain the degree of viral influence on bacterial perturbations observed in IBD related to dysbiosis such as “piggyback” or kill the winner dynamics^43^ with carefully calculated correlations^44^.

Based on the viral proteomes it is clear that a majority of HQ viruses extracted in the bulk metagenomes are likely temperate as we have found integrase proteins in 46% of the viral populations and Lambda-repressor proteins in 60% of viruses infecting *Faecalibacterium* bacteria. This adds to the expectation that the non-enriched viromes can be biased toward viruses that infect the dominant host cells in the sample^19^. Interestingly, we found examples of viruses encoding proteins with immunodominant epitopes such as the TonB plug domain (PF07715) and TonB-dependent beta-barrel (PF00593)^42^ in hundreds of viral proteomes extracted from viruses infecting members of *Bacteroidetes* such as *Bacteroides* and *Alistipes*. A recent study has shown that common structural phage proteins such as tail length tape measure protein (TMP) also harbour immunodominant epitopes with influence on anti-tumour immunity^45^. It is therefore interesting to investigate the extent to which viral organisms can influence the human host-microbiota immune balance through horizontal transfer and expression of immunogenic proteins.

Metavirome studies have until now been the primary source for exploring viral diversity in microbiomes. Now, viral populations are increasingly uncovered in bulk metagenomes. Our approach allowed precise clustering of both viral and bacterial populations in three cohorts that enabled direct investigation into viral-host interactions and discovery of new diversity. We believe that future studies can greatly leverage this approach to conduct virome analysis and investigate the viral influence of the intricate microbiome ecosystem that governs human health.

## Supplementary Figures

Supplementary Figure 1. Virome QC evaluation of metaviromes.

Supplementary Figure 2. Virsorter2 prediction statistics.

Supplementary Figure 3. Jumbo viruses in HMP2.

Supplementary Figure 4. Viral taxonomy percentages for datasets.

Supplementary Figure 5. Average nucleotide identity (ANI) distributions.

Supplementary Figure 6. Phylogenetic tree of crAss-like viruses.

Supplementary Figure 7. Principal component analysis (PCoA) based on MAGs in HMP2

Supplementary Figure 8. Bacterial alpha diversity metrics HMP2.

Supplementary Figure 9. Abundance patterns of associated Viruses and MAGs

Supplementary Figure 10. MAGs connected to Viruses by Phylum in HMP2.

Supplementary Figure 11. Viral-host prediction benchmark on the HMP2 dataset.

Supplementary Figure 12. Diversity generating region (DGR) specificity.

Supplementary Figure 13. Virsorter2 predictions for Viruses and Dark-matter groups in HMP2.

## Supplementary Tables

Supplementary Table 1. Random Forest (RF) performance table.

Supplementary Table 2. Overcomplete genomes VAMB bins vs single-contig evaluation.

Supplementary Table 3. Viral taxonomy counts for datasets.

Supplementary Table 4 (excel file). Counts of viral proteins.

Supplementary Table 5 (excel file). Enriched Viral proteins by predicted Host taxonomy.

## Methods

### Datasets

The Copenhagen Prospective Studies on Asthma in Childhood 2010 (COPSAC) dataset consisted of 662 paired samples obtained at age 1 year from an unselected childhood cohort (^32 33^). The Diabimmune dataset contained 112 paired samples from controls and type 1 diabetes patients. The Human Microbiome Project 2 cohort consisting of 1317 metagenomic samples were downloaded from https://ibdmdb.org/tunnel/public/summary.html.

### Processing of metagenomics and metaviromics datasets

Metagenomic samples of infants *en route* T1D recruited to the Diabimmune study were downloaded from https://pubs.broadinstitute.org/diabimmune (October 2019). Metagenomic samples were quality-controlled and trimmed for adaptors using kneaddata (https://github.com/biobakery/kneaddata) and trimmomatic (v.0.36)^46^ settings: ILLUMINACLIP: NexteraPE-PE.fa:2:30:10 LEADING:20 TRAILING:20 SLIDINGWINDOW:4:20 MINLEN:100. Each metagenomic sample was assembled individually using metaspades (v. 3.9.0)^47^ using the parameters ‘--meta, -k 21,33,55,77,99’ and filtered for contigs with minimum length of 2,000 base pairs. Mapping of reads to contigs was done using minimap2 (v.2.6)^48^ using ‘-N 50’ and filtered with samtools (v.1.9)^49^ using ‘-F 3584’. Contig abundances were calculated using jgi_summarize_bam_contig_depths from MetaBAT2 (v.2.10.2)^50^. Metagenomic bins were defined using VAMB (v. 3.1)^30^ to cluster the metagenomic contigs into putative MAGs and viruses. Initially, the contents of all bins were searched for viral proteins with hmmsearch (v. 3.2.1)^51^ against VOGdb (v. 95) (https://vogdb.csb.univie.ac.at/). The presence of bacterial hallmark genes were determined using both CheckM (v.1.1.2)^52^ and hmmsearch against the miComplete bacterial marker HMM database (v.1.1.1)^53^. A viral-score of each contig was computed using DeepVirFinder (DVF v.1.0)^26^. We initially assessed the metaviromes of the COPSAC and Diabimmune datasets using ViromeQC^54^ and found 5.1 and 0.21 times viral enrichment of the two datasets, respectively (**Supplementary Figure 1**).

### Training the Random Forest to predict viral bins

First we established an initial viral truth set in the metagenomic assembly for the random forest classification. For each metagenomics bin we computed the fraction of contigs mapping to a set of non-redundant viral sequences (Gold standard) using blastn (v. 2.8.1)^55^ with a minimum sequence identity of 95% and query coverage of 50%. Gold standard viral contigs of the paired metaviromics datasets were provided by the authors of the Diabimmune and COPSAC studies. Metagenomic bins with >= 95% of contigs matching with the above criteria were annotated as Viral bins for training and validation. For annotating bacterial bins, MAGs were identified using CheckM (v.1.1.2). MAGs with a completeness score of 10% or above and contamination <= 30% were added to the training and validations set labelled as bacteria. For training we used COPSAC and validated using the Diabimmune dataset. Thus the model was trained to distinguish confidently labelled bacterial and viral bins produced by VAMB, this provided a RF model highly effective at removing non-viral bins and providing a highly enriched candidate set of viral bins that could be further evaluated using dedicated validation tools. In the RF-model we included features such as bin-size, the number of distinct bacterial hallmark genes, the number of different PVOGs in a bin divided by the number of contigs in the bin, viral-prediction DVF score (median DVF score for a bin) defined by DeepVirFinder. The Random Forest model was implemented in Python using *RandomForestClassifier* (sklearn v. 0.20.1) with 300 estimators and using the square root of the number of features as the number of max features. The model was trained on the COPSAC dataset using 40% of observations for training and 60% for validation. Subsequently ROC/AUC, recall and precision was calculated using the Diabimmune recovered viruses as an evaluation set. The performance of DVF was assessed on contig-level by predicting contigs with p-value < 0.05 and score > 0.5 as viral else bacterial.

### Intersection of viruses in MGX and MVX data

In order to identify the number of viruses assembled and binned in the metagenomic (MGX) datasets we searched the metavirome (MVX) viruses in all-vs-all search and calculated genome-to-genome average nucleotide identity (ANI) and genome coverage as aligned fraction (AF). Here we defined species level above 95% ANI and strain-level above 97% ANI. Overlapping or also described as highly-similar viruses between the paired MGX and MVX datasets were those fulfilling the ANI>95% & >75% AF criteria. This search was conducted using FastANI (v.1.1, ‘-fragmenlen 500 -minimumfrag 2 -minimum 80% ANI’)^56^ with genome coverage >=50% (bidirectional fragments / total fragments). We note that hits with less than 80% ANI were not included. We expected that we might be able to find fragmented/incomplete viruses assembled in the metavirome but were more curious about near-complete viruses, thus we quality controlled all MVX viruses using CheckV (v0.4.0, default settings, database v. 0.6)^29^ to achieve a completeness estimate for each. By labelling the quality of each MVX virus we organised the success of genome recovery into the 4 CheckV levels (Low-quality <= 50%, Medium-quality >=50%, High-quality >=90%, Complete =100%). Complete = Closed genomes based on direct terminal repeats (DTR), inverted terminal repeats (ITR). Furthermore, we also quality controlled the putative viruses assembled and binned in the MGX to ask the reverse question, i.e. to what extent do we find complete viruses with no similarity to viruses in the MVX.

### Completeness of viruses recovered in metavirome and bulk metagenomes

To standardise our viral recovery performance across different datasets, we used the guidelines on Minimum Information about an Uncultivated Virus Genome (MIUViG)^19^. The viral completeness of viruses from metaviromics data was assigned using CheckV described as above. CheckV was used to conduct a benchmark on virus genome completeness by evaluating single-contig assemblies against the use of viral bins (also described as viral MAGs). To this end, we based our analysis solely on AAI-model predictions. As the authors of CheckV note, the programme is not designed by default to accommodate viral MAGs and may not deal properly with contaminants from bacterial or viral sources^29^. This became clear as we observed a majority of HMM-model predicted viruses consisting of sequences with close to zero percent viral sequence (**Supplementary Figure 2**). We suspect that this was to be expected since the HMM-model is designed for single viral assemblies. I.e. the model cannot deal properly with cases where a viral marker gene is identified in a bin and contaminating sequences inflate the total bin size to randomly fit into the reference size range of viruses encoding the same viral marker. Hence to avoid including false-positive viral bins, we defined a viral population as HQ-ref when at least one bin in the VAMB cluster contained a HQ evaluation based on AAI-evaluation. All viral bins with a CheckV computed genome copy number >= 1.25 were removed to control for ‘concatemers’. Finally, viral bins with an estimated completeness >120% (overcomplete-genomes) were removed as well to control for contaminated bins. We found that the frequency of HQ genomes, which according to MIUViG standards^19,20^ were “overcomplete-genomes” (estimated completeness >120%), was between 7.9-14.2% for the viral bins and 3.8-6.1% for single contig evaluation **(Supplementary Table 2)**. Hence, the binning approach generates more overcomplete-genomes, although these can be identified and removed using for instance CheckV. The remaining populations without a single HQ or MQ bin within their VAMB-cluster were described as dark-matter. For identifying viruses in ‘dark-matter’ populations, we ran Virsorter2 (v.2.0)^57^ and considered sequences or bins with a prediction score >0.75, at least one viral hallmark and a minimum size of 10 kbp as a putative virus. In this subset of putative viruses we defined “viral-like” dark-matter when they had a CRISPR-spacer with a bacterial MAG (see ‘Viral host prediction’).

### Viral taxonomy and function

While the databases of viral genomes continue to grow, taxonomy is still a challenge for viral genomes with little similarity to the International Committee on Taxonomy of Viruses (ICTV) annotated genomes. Viral proteins were predicted using prodigal (v.2.6.3)^58^ using ‘-meta’. All proteins were annotated using viral protein specific databases such as VOG (http://vogdb.org) or viral subsets of TrEMBL used in the tool Demovir (https://github.com/feargalr/Demovir). Viral taxonomy was assigned to each bin using the plurality rule described before in Roux et al (ref. ^20^): (1) taxonomy was assigned to genomes with at least two PVOG proteins using a majority vote (>=50% else NA) on each taxonomic rank based on the last common ancestor (LCA) annotation from the PVOG entries. (2) The CheckV VOGclade taxonomy was transferred if available from the best viral genome match in the CheckV database. In order to annotate “crAss-like” viruses, predicted proteins were aligned using blastp (v. 2.8.1)^55^ to the large subunit terminase (Terl) protein and DNA polymerase (accessions: YP_009052554.1 and YP_009052497.1) of the progenitor-crassphage using already described cutoffs^59^. When investigating taxonomic annotations, considering only MQ-Complete viral bins, the most dominant viral family annotated was Siphoviridae accounting for 53.5% of the viral bins (**Supplementary Figure 4**). Furthermore, we could assign Myoviridae 14.57%, Podoviridae 8.59%, Microviridae 8.30%, crAss-like 3.61%, CRESS 2.52%, Herelleviridae 1.37%, and Inoviridae 0.58%. Finally 6.93% of viruses could not be confidently assigned any viral taxonomy. Similar distributions of taxonomic annotations were also observed for Diabmmune and COPSAC (**Supplementary Table 3**).

For viral proteomes, we utilised CheckV’s contamination detection workflow to extract proteins encoded only in viral-regions to avoid host contamination. These viral proteins were analysed with interproscan (v. 5.36-75.0)^60^ using the following databases: PFAM, TIGRFAM, GENE3D, SUPERFAMILY and GO-annotation. For each annotated functional domain in viruses predicted to infect a given host genus enriched proteins were identified using Fisher’s exact test using the function *phyper* in base R. P-values were adjusted using False discovery rate (FDR) correction^61^. Viral reverse transcriptase enzymes were grouped into DGR-clades by querying each protein sequence against a database of RT DGR clade HMM models while DGR target genes were identified using the methods and pipeline provided^62^.

### Phylogenetic tree of crAss-like viruses

A phylogenetic tree was constructed for crAss-like viruses identified in the HMP2 dataset based on proteins annotated as the large terminase subunit protein (the TerL gene). First, viral bins annotated as “crAss-like” were determined as described above. “crAss-like” proteomes were aligned to a terminase large subunit protein (accession: YP_009052554.1) and also against VOGdb hmmsearch (v. 3.2.1, hmmscore >= 30)^51^ against VOGdb (v. 95) (https://vogdb.csb.univie.ac.at/). The VOG entries corresponding to the terminase large subunit:VOG00419, VOG00699, VOG00709, VOG00731, VOG00732, VOG01032, VOG01094, VOG01180, VOG01426, were identified using a bash command on a VOGdb file: “grep -i terminase vog.annotations.tsv”. An alignment file was produced for proteins, annotated as terminase large subunit, using MAFFT (v. 7.453)^63^ and Trimal (v. 1.4.1)^64^ and converted into a phylogenetic tree using IQtree (v. 1.6.8 -m VT+F+G4 -nt 14 -bb 1000-bnni)^65^.

### Viral host prediction

Viral genomes were connected to hosts using a combination of CRISPR spacers and sequence similarity between viruses categorised as HQ-ref and MAGs. CRISPR arrays were mined from COPSAC and HMP2 MAGs using CrisprCasTyper (v.1.2.3)^66^ with ‘--prodigal meta’ and all spacers were blasted with blastn-short (v. 2.8.1)^55^ against all viral genomes to identify protospacers. CRISPR spacer matches with >=95% sequence identity over 95% of spacer length and maximum 2 mismatches were kept. In order to identify the host of viruses, viral bins were aligned to MAGs using FastANI (v.1.1, ‘--fragLen 5000 --minFrag 1’)^56^ and blastn megablast (v. 2.8.1)^55^ with a minimum ANI >= 90% and sequence identity >= 90, respectively. We followed the approach described by Nayfach et al. (ref. ^40^) to calculate host-prediction consensus and accuracy. The viral host was defined using a plurality rule at each taxonomic rank based on the lineage of bacteria connected using either CRISPR-spacer or alignment to the given virus. The cutoffs described above were selected after benchmarking the alignment approach with FastANI and blastn at various thresholds. We observed an increased host-prediction consensus and accuracy at the species rank using the threshold described above with FastANI with ANI >=90% based on at least one 5000 bp fragment, compared to blastn thresholds described by Nayfach et al. (ref. ^40^). We evaluated the agreement of our two host prediction methods and found up to 58% consensus on host taxonomy on species rank (**Supplementary Figure 11A)**. We further benchmarked host-prediction purity by calculating the most common host for each viral population according to (1) CRISPR-spacer and (2) alignment independently.

Viruses were annotated as a temperate virus if (1) the virus was found to be integrated into a MAG with >=80% query coverage and ANI >=90% or (2) an integrase protein-annotation could be found in the viral proteome. Integrase proteins were determined searching for *integrase* in the InterPro entry description of each interproscan protein annotation (see Viral taxonomy and function for details).

### Differential abundance of viral populations and MAGs

Sample abundance of each viral population was calculated as a mean read per kilobase million (RPKM) of all contigs with at least 75% coverage belonging to a VAMB cluster. Differential abundance analysis of all viruses were tested using the Linear mixed-effect model R-function *lmer* (lme4 package v. 1.1-26)^67^. The model used was ‘Virus ~ dysbiosis_index + diagnosis + sex + (1|Subject)’. Subjects were included as random effects to account for the correlations in the repeated measures (denoted as (1 | subject)) and the log transformed relative abundance of each virus was modelled as a function of diagnosis (a categorical variable with non-IBD as the reference group) and the dysbiosis index (continous covariate) while adjusting for subjects age as a continuous covariate and sex as a binary variable.

## Acknowledgements

We thank Mani Arumugam, Eduardo Rocha, Nicolas Rascovan, Ramnik Xavier and Hera Vlamakis for fruitful discussions. J.J., J.N.N. and S.R. were supported by the Novo Nordisk Foundation (grant NNF14CC0001).

## Author contributions

S.R. conceived the study and guided the analysis. J.J. wrote the software and performed the analyses. S.A.S and J.S, L.D and D.S.N generated metavirome data and created the viral gold standard for COPSAC data. S.S. and J.S. generated COPSAC metagenome data. D.P. and J.N.N. guided the analyses. J.J., S.R. and D.P. wrote the manuscript with contributions from all co-authors. All authors read and approved the manuscript.

## Data availability

The Diabimmune dataset and HMP2 datasets are available from the European Nucleotide Archive with the accessions PRJNA398089 and PRJNA387903. The COPSAC metagenomics data will be available upon publication from the COPSAC consortium. Gold standard virus genomes for COPSAC and Diabimmune were provided by Shiraz Shah and Tommi Vatanen, respectively.

## Code availability

The VAMB code is available at https://github.com/RasmussenLab/vamb and the workflows used in this analysis is available at https://github.com/RasmussenLab/phamb.

