## Supplementary Figures for "Genome binning of viral entities from bulk metagenomics data"

#### Author List Footnotes

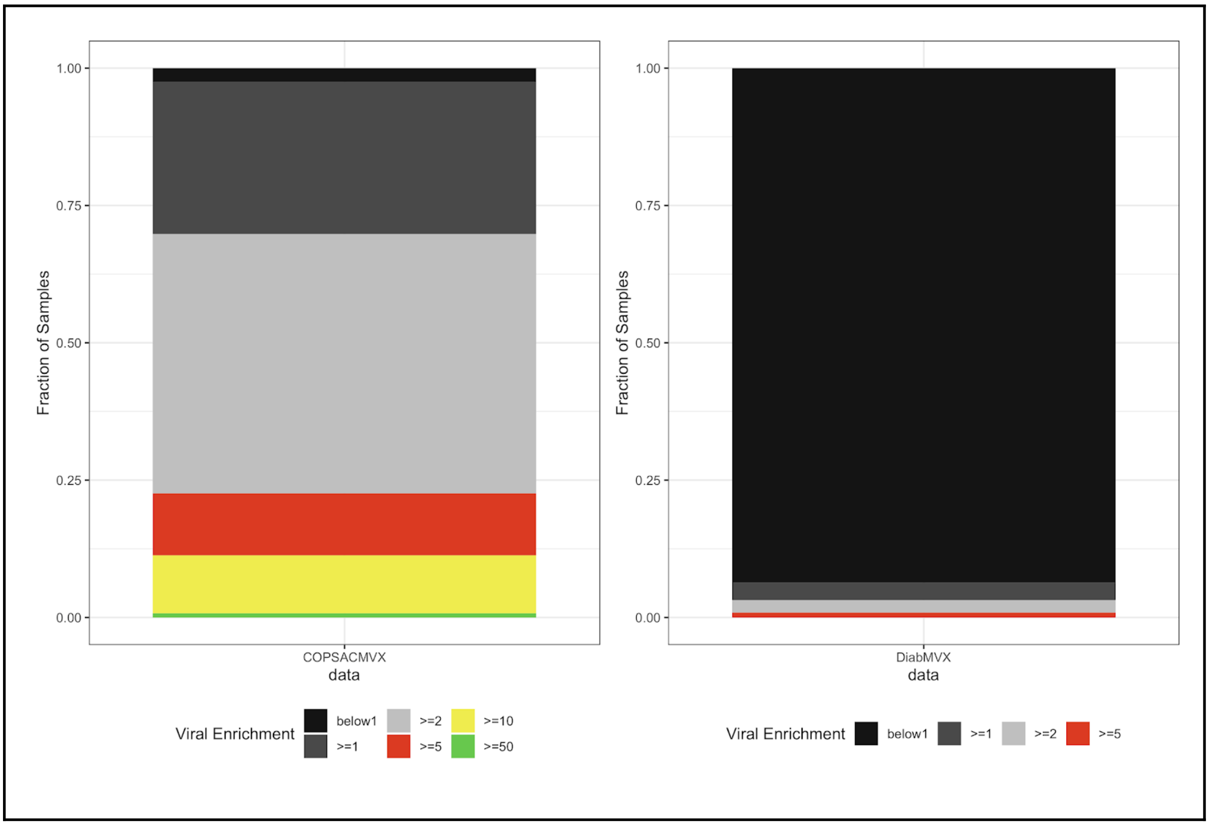


**Supplementary Figure 1. Virome QC evaluation of metaviromes.** All samples in COPSAC and the Diabimmune T1D metavirome datasets were processed with ViromeQC to achieve viral enrichment estimates. The magnitude of enrichment relative to bulk metagenomics is coloured using breaks described in the paper by Zolfo *et al.* where black indicates low and green indicates high enrichment.

Virsorter2 on Diabimmune T1D

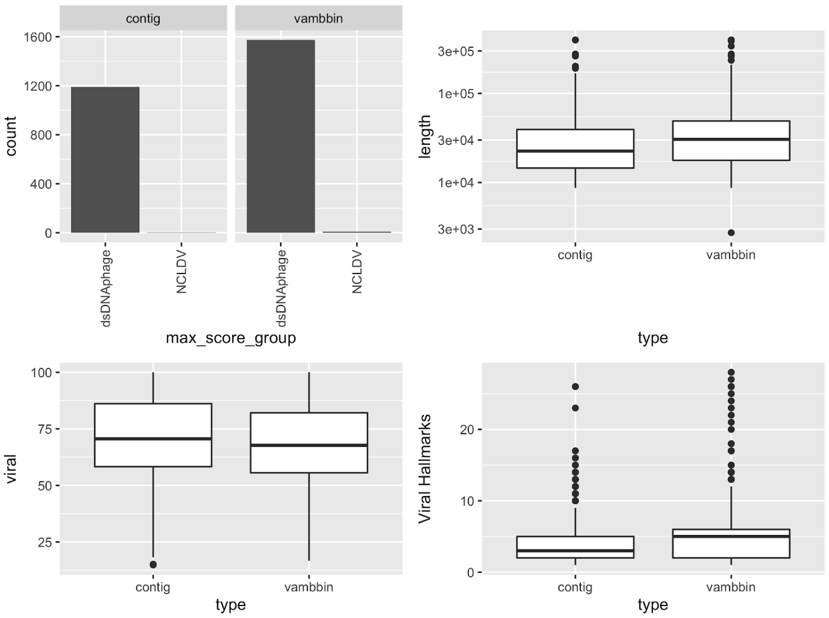


Virsorter2 on COPSAC dataset

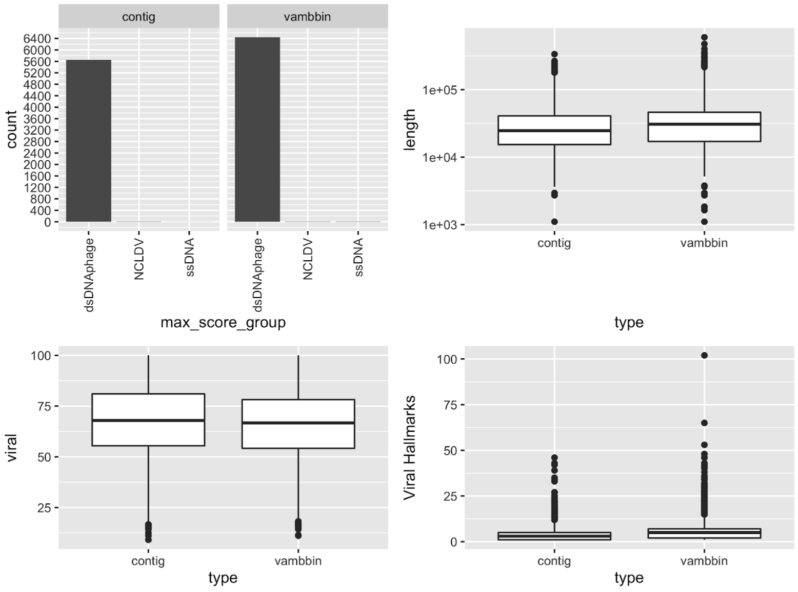


**Supplementary Figure 2. Virsorter2 prediction statistics.** Virsorter2 was run on sequences assembled and binned from COPSAC and Diabimmune bulk metagenomes. Prior to Virsorter2 analysis, sequences were cleaned using CheckV that removes bacterial regions. Results are only shown for sequences predicted as double or single-stranded DNA phage or nucleocytoplasmic large DNA viruses (NCLDV) with a prediction score >0.75 and >=1 viral hallmark gene.

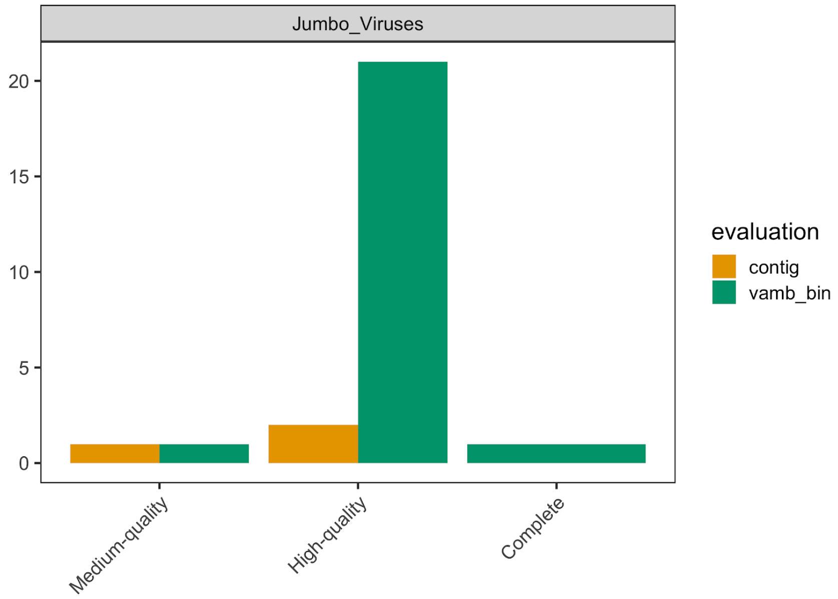


**Supplementary Figure 3. Jumbo viruses in HMP2.** Viral completeness was estimated for both single-contigs and VAMB-bins in HMP2. Evaluation of genome completeness was determined using CheckV here shown for Medium-quality >=50% , High-quality >=90%, Complete =100%). Closed genomes are annotated as "Complete" based on direct terminal repeats or inverted terminal repeats. After adjusting for possible host-contaminating sequences in both single-contigs and VAMB bins, several Jumbo viruses with a size >=200 kbp were found.

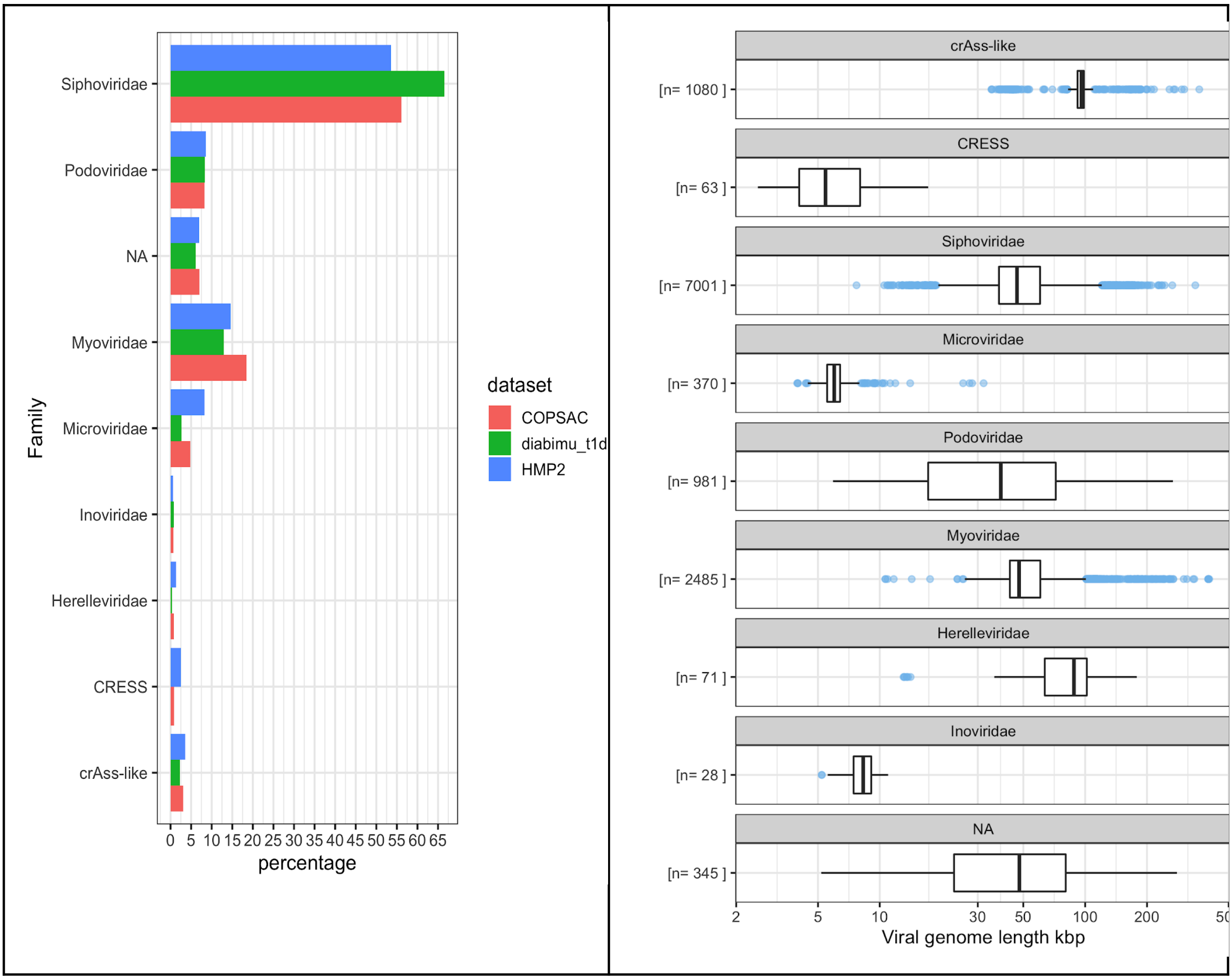


**Supplementary Figure 4. Viral taxonomy percentages for datasets.** Viral taxonomy was assigned to each bin using the plurality rule described before in Roux et al.: (1) taxonomy was assigned to bins with at least two PVOG proteins (VOGdb) using a majority vote (>=50% else NA) on each taxonomic rank based on the last common ancestor (LCA) annotation from the PVOG entries. (2) The CheckV VOGclade taxonomy was transferred if available from the best viral genome match in the CheckV database. CrAss-like viruses were annotated as described by Guerin *et al.* By far, the most frequently annotated viral family was *Siphoviridae* in all datasets.

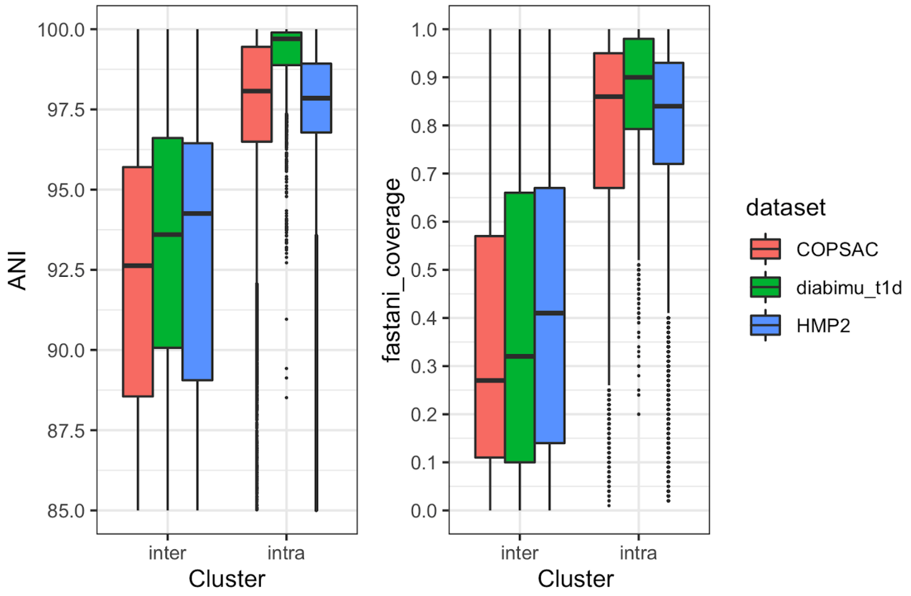


**Supplementary Figure 5. Average nucleotide identity (ANI) distributions.** In order to assess how VAMB clusters highly similar viral bins, we compared ANI between viral bins of different VAMB clusters (inter) and within the same cluster (intra). Coverage was also calculated in a similar way between bins as the number of bidirectional fragments / total fragments. This showed that highly similar viral bins (ANI > 97.5) were consistently clustered into the same VAMB clusters.

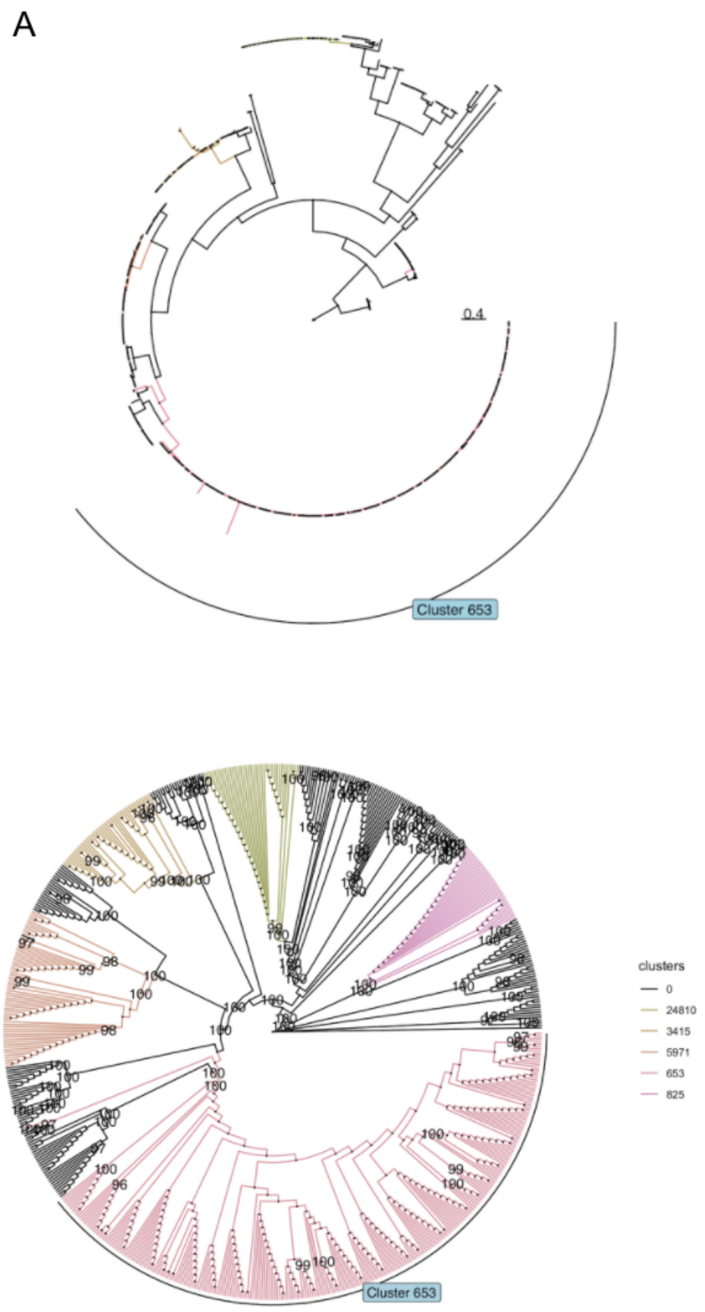


**Supplementary Figure 6. Phylogenetic tree of crAss-like viruses. (a)** A phylogenetic tree was constructed for crAss-like viral bins identified in the HMP2 dataset based on proteins annotated as the large terminase subunit protein (the *terL* gene). Branches drawn according to phylogenetic distance with cluster 653 indicate the progenitor-crassphage. **(b)** The cladogram of the tree, same as Figure 3d, but displayed with bootstrap values calculated by IQtree.

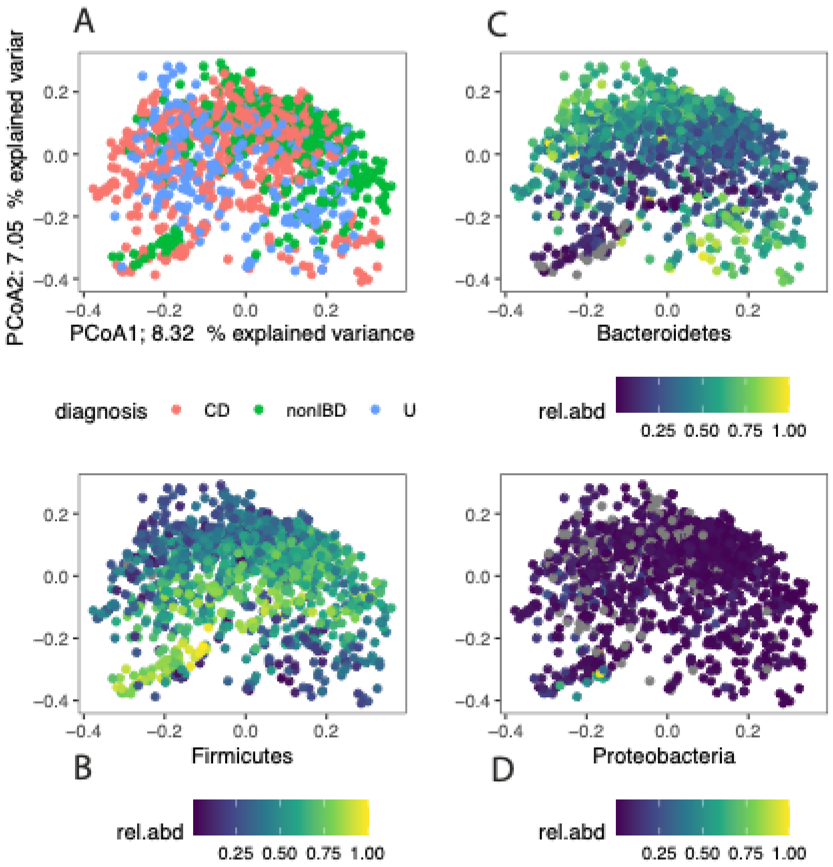


**Supplementary Figure 7. Principal component analysis (PCoA) based on MAGs in HMP2. (a)** PCoA was performed on Bray-Curtis distance matrix of MAG abundances in the HMP2 cohort. Samples were not easily separated based on diagnosis status from the HMP2 metadata **(b-d)** Instead, differences in the proportion of relative abundance occupied by Bacteroidetes and Firmicutes bacteria was reflected by the PCoA analysis.

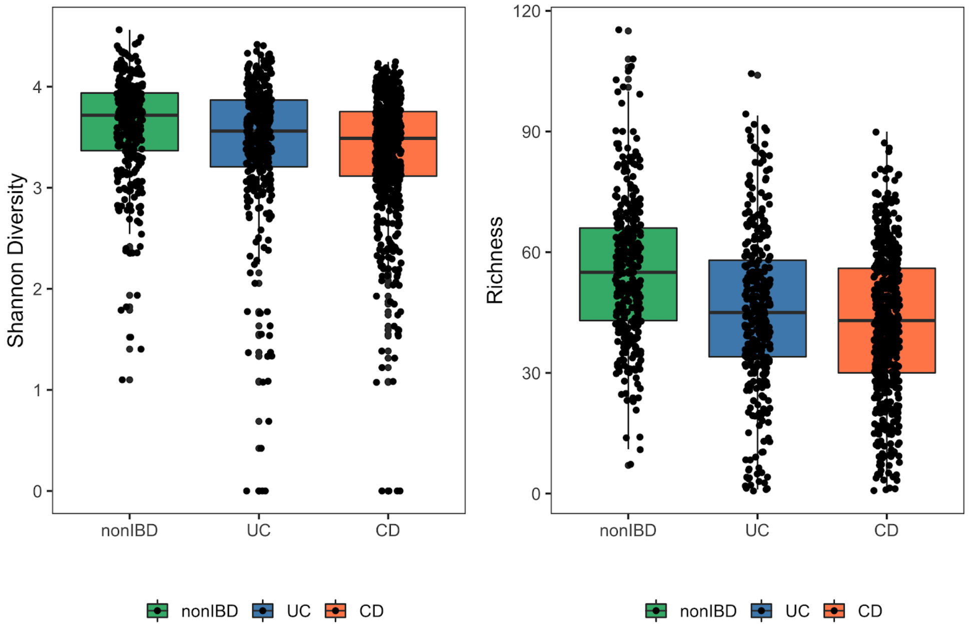


**Supplementary Figure 8. Bacterial alpha diversity metrics of the HMP2.** The bacterial composition of HMP2 samples were characterised with alpha-diversity metrics such as Shannon Diversity index (vegan package in R) and Richness (the number of different MAGs with abundance above zero). A general downward shift in both metrics was observed for samples from subjects with diagnosed ulcerative colitis (UC) and Crohn's disease (CD) relative to samples from nonIBD (control) subjects.

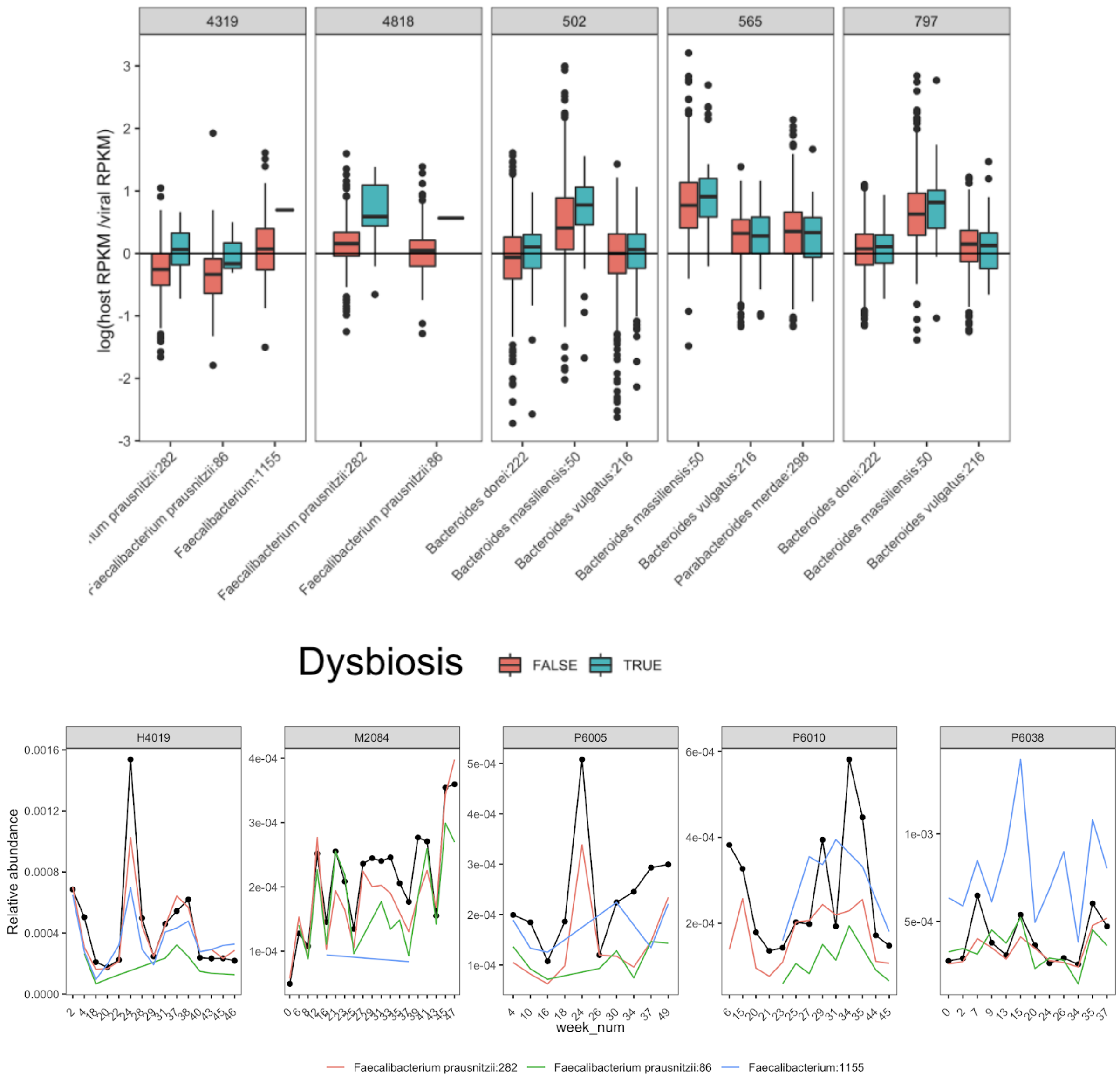


**Supplementary Figure 9. Abundance patterns of associated Viruses and MAGs. (Upper panels)** Viral-bacterial relationships were determined using a combination of CRISPR-spacer and sequence alignment. In order to confirm host-dependencies for putative temperate viruses (each panel), the ratio of relative abundance was calculated across samples for a given virus and bacterial host(s). Furthermore, these abundance ratios were illustrated for dysbiotic samples to capture potential shifts or breaks in host-dependency. **(Lower panels)** The relative abundance of individual viruses (viral *cluster 4319* indicated with the black line) can be visualised along associated bacterial hosts (coloured lines) over time in different subjects (each panel).

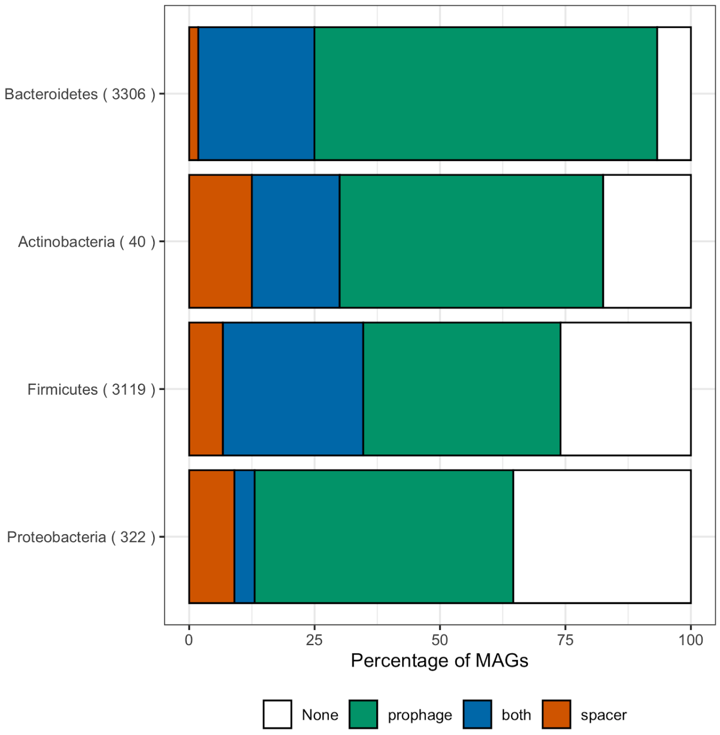


**Supplementary Figure 10. MAGs connected to Viruses by Phylum in HMP2.** Host-viral associations were determined using CRISPR spacers, which were mined from MAG bins, and viral bin alignments to MAG bins. The percentage of bacterial bins annotated with a viral population is illustrated below on phylum rank. Here the majority of Bacteroidetes bins are only annotated to a virus through viral alignment as *prophage* evidence (green), second is where *both* (blue) a viral alignment and CRISPR spacer match the given virus. Finally, some bacterial bins only have CRISPR *spacers* (red) against viruses and no viral alignment.

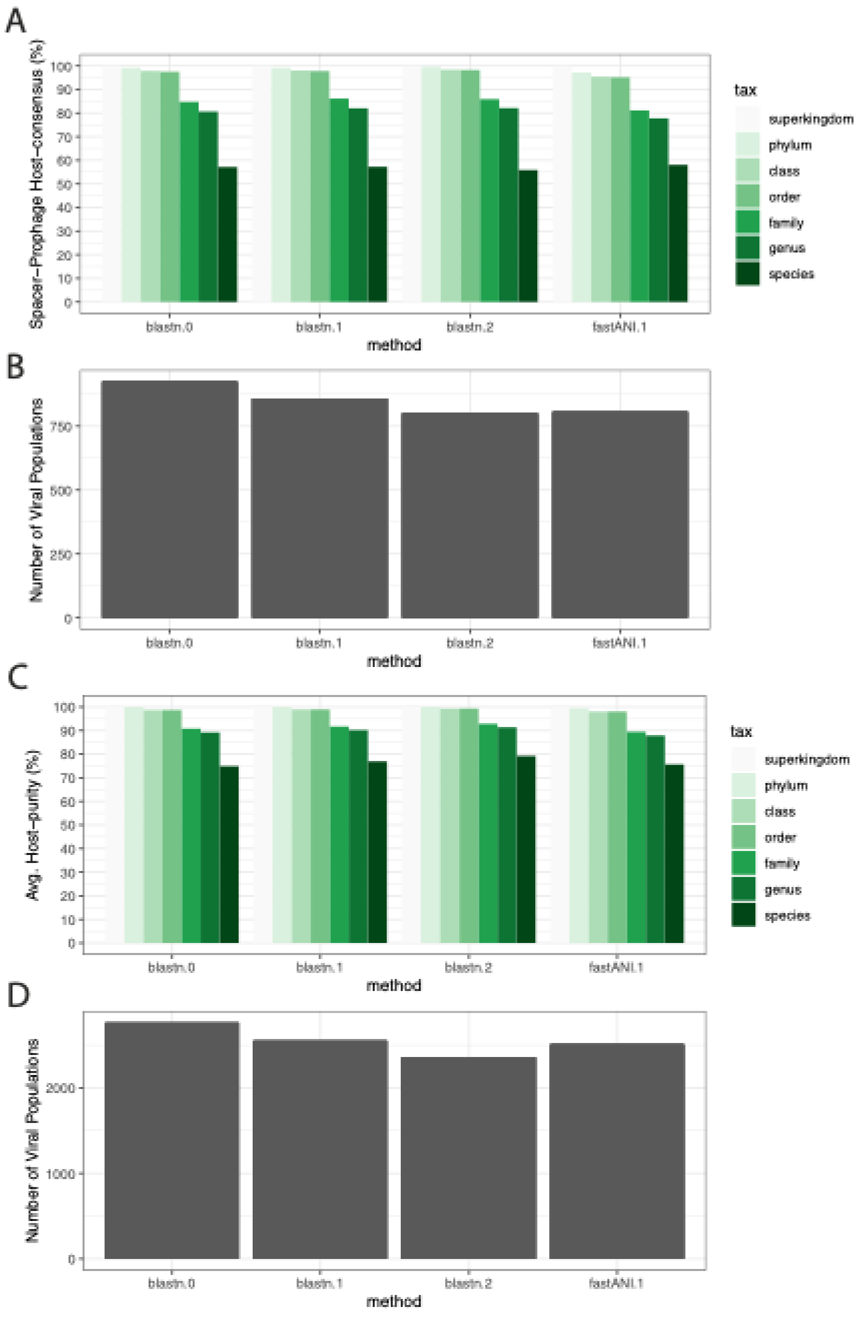


**Supplementary Figure 11. Viral-host prediction benchmark on the HMP2 dataset.**

**(a).** For viruses with hosts predicted by both methods (genome-alignment and CRISPR-spacers), the average host-prediction consensus was calculated across multiple taxonomic ranks. The consensus was calculated using three different cutoffs with blastn and one cutoff for FastANI. In addition, the number of viral populations host annotated at each threshold is shown in **(b)**. We also calculated a “purity” measurement of predicted hosts for each virus in **(c)**. I.e. if a virus genome aligns to three different MAGs and MAG taxonomy on species rank is [B. vulgatus, B. vulgatus, B. dorei] the virus host purity is 66% on species rank. For this benchmark the number of viral populations used is displayed in **(d).**

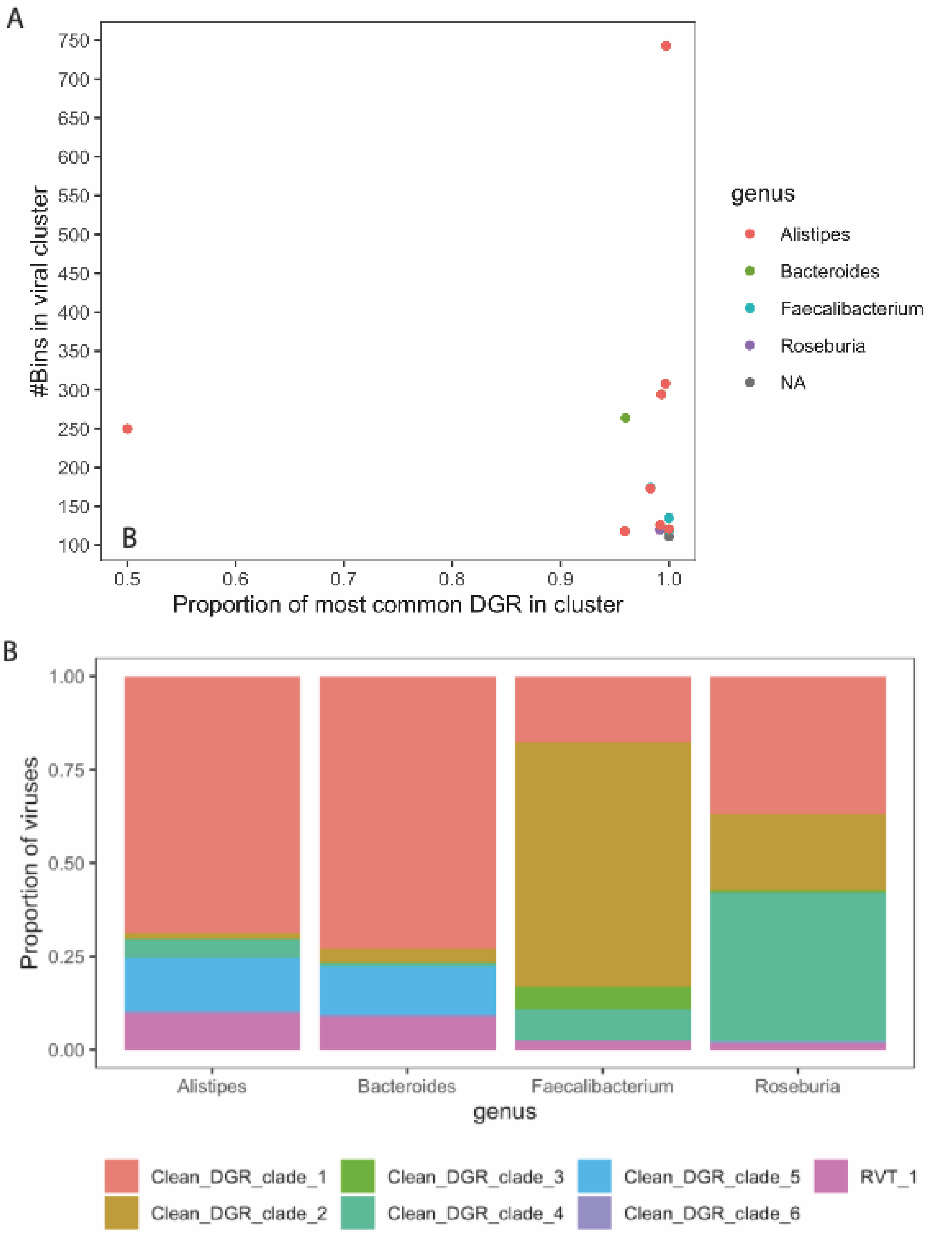


**Supplementary Figure 12. Diversity generating region (DGR) specificity. (a)** DGR specificity, the most common DGR type within a VAMB cluster, for viral populations coloured by predicted host taxonomy on genus rank. The DGR-clade for reverse transcriptase (RT) proteins were characterised using methods described in Roux et al., 2020. The most frequent DGR was calculated for each viral population to determine DGR specificity. **(b)** DGR-specificity by viral host taxonomy on genus rank.

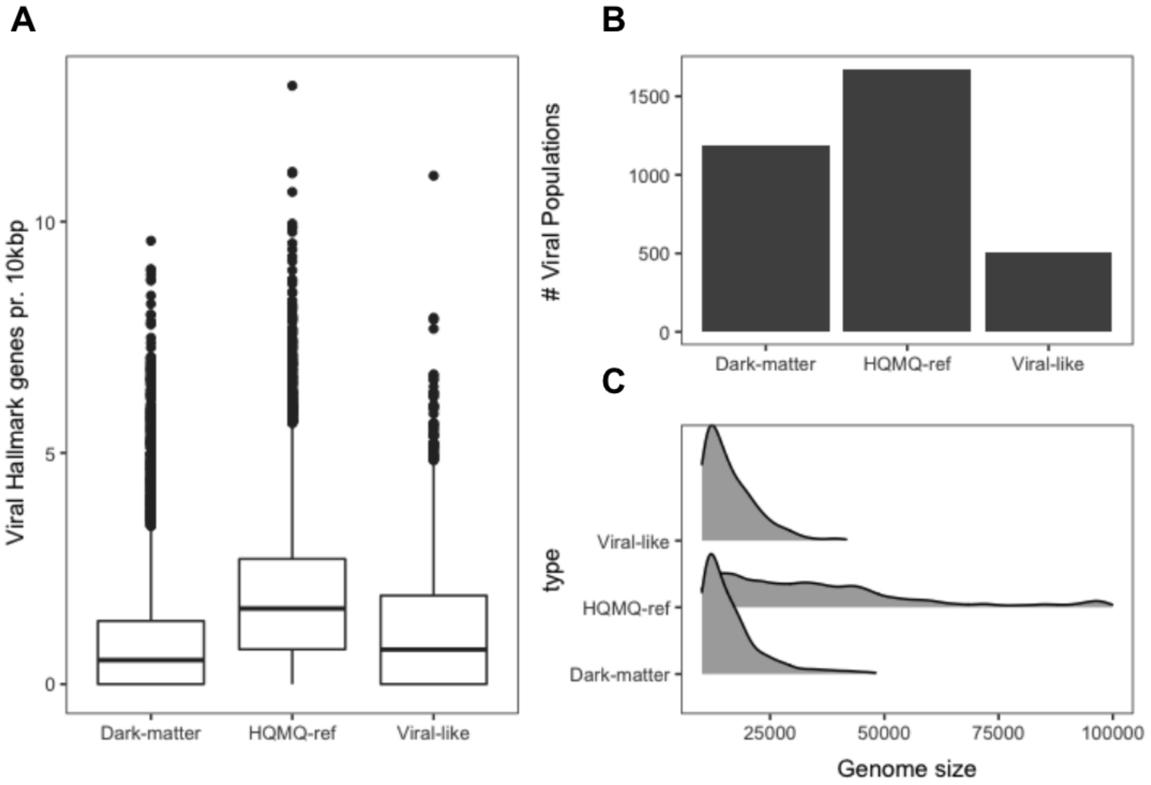


**Supplementary Figure 13. Virsorter 2 predictions for Viruses and Dark-matter groups in HMP2.** VAMB clusters/populations were defined as HQMQ-ref with a high-quality or medium-quality viral bin, else as Dark-matter. Dark-matter populations with CRISPR-spacers against a bacterial MAG were annotated as Viral-like. All bins within each group with a minimum size of 10 kbp were analysed using Virsorter revealing high prediction prediction scores for bins found in all groups (A). **(b-c)** The number of viral hallmarks was higher in the HQMQ group which was also the group with the highest number of bins with a score >0.75 (C). **(d)** In addition, the HQMQ group also comprises larger viruses, suggesting that Dark-matter and Viral-like contains smaller or fragmented viruses.

### Supplementary Tables

| Model name | AUC | Recall | Precision | MCC | F1 |
|---|---|---|---|---|---|
| **Default RF model** | **0,999** | **0,998** | **0,998** | **0,997** | **0,999** |
| **Best random model** | **0,999** | **0,999** | **0,998** | **0,998** | **0,999** |
| **Best grid model** | **0,999** | **0,999** | **0,998** | **0,998** | **0,999** |

**Supplementary Table 1. Random Forest (RF) performance table.** The RF models were built and trained as described in Methods. Validation performance metrics such as area under curve (AUC), recall, precision, Matthews correlation coefficient (MCC) and F1 (2*precision*recall/precision+recall) were calculated based on the Diabimmune dataset.

| Dataset | Evaluation | Overcompleteness | Count | Percentage |
|---|---|---|---|---|
| **Diabimmune** | **contig** | **High-quality** | **151** | **96,17** |
| **Diabimmune** | **contig** | **Unsure (completeness > 120%)** | **6** | **3,82** |
| **Diabimmune** | **vamb_bin** | **High-quality** | **326** | **85,79** |
| **Diabimmune** | **vamb_bin** | **Unsure (completeness > 120%)** | **54** | **14,21** |
| **COPSAC** | **contig** | **High-quality** | **1080** | **95,83** |
| **COPSAC** | **contig** | **Unsure (completeness > 120%)** | **47** | **4,17** |
| **COPSAC** | **vamb_bin** | **High-quality** | **1812** | **87,88** |
| **COPSAC** | **vamb_bin** | **Unsure (completeness > 120%)** | **250** | **12,12** |
| **HMP2** | **contig** | **High-quality** | **2272** | **93,92** |
| **HMP2** | **contig** | **Unsure (completeness > 120%)** | **147** | **6,08** |
| **HMP2** | **vamb_bin** | **High-quality** | **6127** | **92,08** |
| **HMP2** | **vamb_bin** | **Unsure (completeness > 120%)** | **527** | **7,92** |

**Supplementary Table 2. Overcomplete genomes VAMB bins vs single-contig evaluation.** We investigated the frequency of viral bins with a viral size quite greater than the anticipated reference virus determined by the CheckV AAI-model. We tallied the number of High-quality bins determined and counted the ones with completeness > 120% for all datasets. This was also done for HQ single-contigs evaluated using the CheckV AAI-model.

| Dataset | Family | # Genomes | # Populations | % of populations |
|---|---|---|---|---|
| COPSAC | crAss-like | 143 | 38 | 3,11 |
| COPSAC | CRESS | 12 | 10 | 0,81 |
| COPSAC | Herelleviridae | 18 | 9 | 0,73 |
| COPSAC | Inoviridae | 15 | 8 | 0,65 |
| COPSAC | Microviridae | 71 | 59 | 4,83 |
| COPSAC | Myoviridae | 496 | 225 | 18,42 |
| COPSAC | NA | 99 | 86 | 7,04 |
| COPSAC | Podoviridae | 278 | 101 | 8,27 |
| COPSAC | Siphoviridae | 1536 | 685 | 56,10 |
| Diabmmune | crAss-like | 21 | 6 | 2,28 |
| Diabmmune | Herelleviridae | 1 | 1 | 0,38 |
| Diabmmune | Inoviridae | 2 | 2 | 0,76 |
| Diabmmune | Microviridae | 10 | 7 | 2,66 |
| Diabmmune | Myoviridae | 74 | 34 | 12,92 |
| Diabmmune | NA | 25 | 16 | 6,08 |
| Diabmmune | Podoviridae | 38 | 22 | 8,34 |
| Diabmmune | Siphoviridae | 356 | 175 | 66,54 |
| HMP2 | crAss-like | 916 | 50 | 3,61 |
| HMP2 | CRESS | 51 | 35 | 2,52 |
| HMP2 | Herelleviridae | 52 | 19 | 1,37 |
| HMP2 | Inoviridae | 11 | 8 | 0,58 |
| HMP2 | Microviridae | 289 | 115 | 8,30 |
| HMP2 | Myoviridae | 1915 | 202 | 14,57 |
| HMP2 | NA | 221 | 96 | 6,93 |
| HMP2 | Podoviridae | 665 | 119 | 8,59 |
| HMP2 | Siphoviridae | 5109 | 742 | 53,53 |

**Supplementary Table 3. Viral taxonomy counts for datasets.** Viral bins were taxonomically annotated as described in the Methods section *Viral taxonomy and function*. In the table, the percentage of bins annotated to a given viral family is shown. In addition, the number of distinct viral populations annotated is also shown.

**Supplementary Data 4 (excel file). Counts of viral proteins**

**Supplementary Data 5 (excel file). Enriched viral proteins by predicted host taxonomy**
